# Alcohol-induced Hsp90 acetylation is a novel driver of liver sinusoidal endothelial dysfunction and alcoholic liver disease

**DOI:** 10.1101/2020.08.25.267096

**Authors:** Yilin Yang, Panjamaporn Sungwung, Yirang Jung, Reiichiro Kondo, Matthew McConnell, Teruo Utsumi, William C. Sessa, Yasuko Iwakiri

## Abstract

**Background:** It is unknown whether liver sinusoidal endothelial cells (LSECs) metabolize alcohol. Chronic alcohol consumption decreases endothelial nitric oxide synthase (eNOS)-derived NO production typical of LSEC dysfunction. Heat shock protein 90 (Hsp90) interacts with eNOS to increase its activity. Cytochrome P450 2E1 (CYP2E1) is a key enzyme in alcohol metabolism and facilitates protein acetylation via acetyl-CoA, but its expression in LSECs is unknown. This study investigates alcohol metabolism by LSECs, the mechanism of alcohol-induced LSEC dysfunction and a potential therapeutic approach for alcohol-induced liver injury.

**Methods:** Primary human, rat and mouse LSECs were used. Histone deacetylase 6 (HDAC6) was overexpressed specifically in liver ECs using an adeno-associated virus (AAV)-mediated gene delivery system to decrease Hsp90 acetylation in ethanol fed mice.

**Results:** LSECs expressed CYP2E1 and alcohol dehydrogenase 1 (ADH1) and metabolized alcohol. Ethanol induced CYP2E1 in LSECs, but not ADH1. Alcohol metabolism by CYP2E1 increased Hsp90 acetylation and decreased its interaction with eNOS along with a decrease in NO production. A non-acetylation mutant of Hsp90 increased its interaction with eNOS and NO production, whereas a hyper-acetylation mutant decreased NO production, compared with wildtype Hsp90. These results indicate that Hsp90 acetylation is responsible for decreases in its interaction with eNOS and eNOS-derived NO production. Adeno-associated virus 8 (AAV8)-driven HDAC6 overexpression specifically in liver ECs deacetylated Hsp90, restored Hsp90’s interaction with eNOS and ameliorated alcohol-induced liver injury in mice.

**Conclusion:** Restoring LSEC function is important for ameliorating alcohol-induced liver injury. To this end, blocking acetylation of Hsp90 specifically in LSECs via AAV-mediated gene delivery has the potential to be a new therapeutic strategy.

## Introduction

Liver sinusoidal endothelial cells (LSECs) are indispensable for liver homeostasis (1). LSEC dysfunction has been reported in various liver diseases, including alcoholic liver disease (2–5). However, it is not known whether LSECs metabolize alcohol and, if so, how it is related to alcohol-induced liver injury.

Alcohol is metabolized in the liver by several pathways (6, 7). The most common of these involves two enzymes, alcohol dehydrogenase 1 (ADH1) and aldehyde dehydrogenase (ALDH). In this pathway, alcohol is metabolized first by ADH1 to acetaldehyde, a highly toxic metabolite and a known carcinogen (8). Then, ALDH catabolizes acetaldehyde to acetate, a less active product, which can be converted to acetyl-CoA by acetyl-CoA synthetase (ACSS) (8).

Cytochrome P450 2E1 (CYP2E1) is another enzyme that metabolizes alcohol to acetaldehyde (6), especially in pathological conditions. Ethanol increases the rate of CYP2E1 gene transcription, and also stabilizes CYP2E1 by preventing it from degradation (9). CYP2E1 is mainly expressed in the liver with its highest expression in hepatocytes. Accordingly, the study of CYP2E1 in the liver has been concentrated on hepatocytes, although Kupffer cells are also known to express it to some degree (10, 11). The presence and function of CYP2E1 in other hepatic cells, including LSECs, remain unknown.

In this study, we have explored the presence of CYP2E1 and ADH1 as well as alcohol metabolism by these enzymes in LSECs. We have also examined the mechanism of LSEC dysfunction represented by decreased NO production and liver injury caused by CYP2E1-mediated alcohol metabolism as well as the therapeutic strategy of blocking this mechanism using an adeno-associated virus (AAV)-mediated gene delivery system.

Heat shock protein 90 (Hsp90) is a chaperone protein that helps protein homeostasis by assisting other proteins to function (12). Acetylation of Hsp90 reduces the formation of complexes between Hsp90 and its client proteins and thus has a negative impact on their function (13). In endothelial cells, Hsp90 increases eNOS activity by directly binding to eNOS (14, 15), and thereby promotes NO production. Given that acetyl-CoA, a final product of alcohol metabolism, serves as a substrate for protein acetylation (16), we thus hypothesized that alcohol metabolism by CYP2E1 could cause excessive Hsp90 acetylation and decrease its interaction with eNOS, leading to decreased NO production in LSECs and contributing to alcohol-induced liver injury. We also hypothesized that overexpression of histone deacetylase 6 (HDAC6) in LSECs, which deacetylates Hsp90, could restore eNOS activity and NO production, thereby ameliorating liver injury.

In this study, we have determined CYP2E1/ADH1 expression and alcohol metabolism in LSECs for the first time. We have also shown that alcohol metabolism by CYP2E1 increases Hsp90 acetylation in LSECs and decreases its interaction with eNOS, leading to decreased NO production, and that liver EC-specific overexpression of HDAC6 restores the chaperone function of Hsp90 and reduces alcohol-induced liver injury in mice. The study suggests that maintaining NO levels in LSECs, and thus normal LSEC function, is important for protecting the liver from alcohol-induced injury. Blocking Hsp90 acetylation in LSECs could be a therapeutic strategy for the treatment of alcoholic liver disease.

## Materials and Methods

### Animal experiments

To examine the effect of HDAC6 overexpression in liver endothelial cells on alcoholic liver disease, we used Cdh5-CreERT2 mice and the NIAAA model for generation of alcoholic liver disease with slight modifications. Cre recombinase expression in endothelial cells of Cdh5-CreERT2 mice was induced by intraperitoneal injection of tamoxifen (Sigma-Aldrich, St. Louis, MO) at a dose of 100 μg/g body weight for 5 consecutive days. Three weeks after the last tamoxifen injection to ensure Cre recombinase expression, Cdh5-CreERT2 mice were administrated with an adeno-associated virus 8 (AAV8)-EF1a-DIO-mKate2 (Vector Biolabs, Malvern, PA) (control, n=10) or AAV8-EFS-DIO-mHDAC6 vector (Vector Biolabs) (n=11) through the retro-orbital sinus at a dose of 2 × 10_12_ GC/100 μL per mouse. Three weeks later, mice (7 Kate2 vs. 7 HDAC6) started to be fed with a 5% ethanol-liquid diet (F1258SP, Bio-Serv, Flemington, NJ) for 10 days. On the morning of Day 11, they received an oral gavage of ethanol (3.75 g/kg body weight) and were sacrificed 9 hours later to collect blood and liver tissues. For a control group, Cdh5-CreERT2 mice (3 Kate2 vs. 4 HDAC6) were fed with a control liquid diet (F1259SP, Bio-Serv) for 10 days and were sacrificed for sample collection. All mice were approximately 4 months old at the time of sacrifice. We received liver specimens of ethanol fed Sprague Dawley (SD) rats (the NIAAA model) from Dr. Varman Samuel (Yale University).

All animal experiments were approved by the Institutional Animal Care and Use Committee of the Veterans Affairs Connecticut Healthcare System, and performed in adherence with the National Institutes of Health Guidelines for the Use of Laboratory Animals.

### Cell isolation

Primary rat liver sinusoidal endothelial cells (LSECs) and hepatocytes were isolated from SD rats at the Yale Liver Center Core Facility. Briefly, after collagenase perfusion, cell suspension was centrifuged at 50 xg for 3 minutes at room temperature to obtain hepatocytes (pellet fractions) (17). A non-parenchymal cell fraction (NPC, supernatants) was transferred to a new tube, pelleted at 350 xg for 10 minutes and resuspended in EBM-2 Basal Medium (CC-3156, Lonzs, Basel, Switzerland) supplemented with Microvascular Endothelial Cell Growth Medium-2 SingleQuots Supplements (CC-4147, Lonza) and 15% fetal bovine serum. The NPC suspension was subjected to a density gradient centrifugation using Percoll (Lot 10221921, GE Healthcare, Chicago, IL) at 900 xg for 20 minutes at room temperature. An LSEC-enriched fraction was isolated and seeded on collagen-coated coverslips or cell culture dishes. At 3, 24, 48 or 72 hours after cell culture in the endothelial cell growth medium described above at 37oC in a humidified atmosphere containing 5% CO_2_, LSECs were used for immunofluorescence staining or Western blot analysis.

Primary human LSECs were purchased from ScienCell Research Laboratories (#5000, Carlsbad, CA). Human LSECs were cultured in endothelial cell medium (#1001, ScienCell Research Laboratories) supplemented with 5% fetal bovine serum (#0025, ScienCell Research Laboratories), 1% endothelial cell growth supplement (#1052, ScienCell Research Laboratories) and 1% penicillin/streptomycin (#0503, ScienCell Research Laboratories).

We obtained monkey kidney epithelial cells (COS-7) and human hepatocyte cells (HepG2) from ATCC (Manassas, VA), and bovine aortic endothelial cells (BAECs) from Clonetics (San Diego, CA). All cells were grown in Dulbecco’s Modified Eagle Medium (DMEM) (11965092, Gibco, Waltham, MA) supplemented with 10% fetal bovine serum (Gibco) and 1% penicillin/streptomycin (Sigma-Aldrich).

### Hsp90 mutant constructs and eNOS construct

Hsp90α acetylation mutants tagged with FLAG were kind gifts from Dr. Len Neckers (National Cancer Institute). A mutation of the lysine 294 residue to glutamine (Q) represents a hyper-acetylated state of Hsp90, while an arginine (R) mutant represents a non-acetylated state (13). An empty pcDNA3 vector was used as a control, since it was used for the backbone of Hsp90α and eNOS constructs (18).

### *In vitro* ethanol experiment

For ethanol treatment, endothelial cells were seeded on 6-well plates. After treatment with 100 mM ethanol in culture medium for 20 hours, an additional 100 mM ethanol was added into medium and incubated for 4 hours. For trichostatin A (TSA) (203-17561, Wako Chemicals USA, Richmond, VA) or sodium acetate (3470-01, J.T.Baker, Phillipsburg, NJ) treatment, endothelial cells were seeded on 6-well plates and treated with 5 μM TSA or sodium acetate for 24 hours.

### CYP2E1 or ADH1 silencing in human LSECs

Human LSECs were plated on 6-well dishes overnight. For transfection of CYP2E1 siRNA (M-010134-01-0005, Dharmacon, Lafayette, CO), cells were incubated in 500 μl Opti-MEM I Reduced Serum Medium (Life Technologies, Grand Island, NY) containing 20 nM CYP2E1 siRNA or control scrambled siRNA with 3 μl of a transfection agent (ScreenFect A, Wako Chemicals USA) for 6 hours. Then, 500 μl of DMEM was added to each well. After 24 hours, cells were treated with 100 mM ethanol for 24 hours. The same procedures were performed for transfection of ADH1 siRNA (M-006520-01-0005, Dharmacon) and *in vitro* ethanol experiments.

### Transfection

COS-7 cells were seeded on 6-well plates and transfected the next day with 1 μg of WT-eNOS and 1 μg of Hsp90 mutant constructs or pcDNA using Lipofectamine 2000 (Life Technologies) in Opti-MEM I Reduced Serum Medium (Life Technologies) according to the manufacturer's instructions. Cells were used 24 hours after transfection.

### NO assay

Cells were seeded on 6-well plates. Phenol red free culture medium (31053036, Gibco; #1001, ScienCell Research Laboratories) was used during treatment. After treatment, the media was collected and processed for the measurement of nitrite, a stable end product of NO in aqueous solution. Nitrite concentrations were measured by the DAN assay (19).

### Acetyl-CoA assay

Refer to the Supplemental Materials.

### Serum alanine aminotransferase (ALT) measurement

Refer to the Supplemental Materials.

### Histological analysis

Refer to the Supplemental Materials.

### CK8 immunohistochemistry and quantification

Refer to the Supplemental Materials.

### Immunofluorescence

Refer to the Supplemental Materials.

### Quantitative real-time polymerase chain reaction

Refer to the Supplemental Materials.

### Western blot analysis

Refer to the Supplemental Materials.

### Immunoprecipitation

Cells or liver tissues were homogenized in immunoprecipitation (IP) lysis buffer (50 mM Tris-HCl, 150 mM sodium chloride, 1% NP-40 and 0.5% sodium deoxycholate) with protease inhibitor cocktail (Roche Diagnostics, Mannheim, Germany) and kept on ice for 30 minutes. Lysates were centrifuged at 12,000 rpm for 10 minutes at 4°C. Protein concentrations in the supernatants were determined using the BCA assay. For each sample, 750 μg protein was added to 300 μl IP lysis buffer, and the solution was allocated to two centrifuge tubes (200 μl for IP and 100 μl for input). 200 μl IP lysate samples from each group were incubated with primary antibodies such as eNOS (610297, BD Biosciences, San Jose, CA), acetyl-lysine (9441, Cell Signaling Technology, Danvers, MA), Hsp90 (610419, BD Biosciences) and control IgG from same species on a shaker at 4°C. After 24 hours of incubation, protein A-Agarose or protein G-Agarose was added to each sample, followed by additional incubation at 4°C for 3 hours on a rocking platform. Agarose-antibody-antigen complexes were then centrifuged at 500 xg for 30 seconds at 4°C. Agarose pellets were washed with 500 μl IP lysis buffer and centrifugated (500 xg, 30 seconds, 4°C). This process was repeated 3 times. After washing, agarose pellets were resuspended with 60 μl gel-loading buffer and denatured by heating. Each sample was centrifuged, and 15 μl of the supernatant was loaded for Western blot analysis.

### Statistical analysis

Data were expressed as mean ± standard error of mean (SEM). Statistical significance was determined by performing one-way analysis of variance (ANOVA) or Student’s t-test. A *p*-value < 0.05 was considered statistically significant.

## Results

### CYP2E1 is expressed in rat, mouse and human LSECs and upregulated in response to ethanol

We observed that cytochrome P450 2E1 (CYP2E1) was expressed in rat primary LSECs (Figs. 1A, B&C), while hepatocytes, the major CYP2E1-expressing cells in the liver, exhibited approximately 4.5-fold higher protein levels (*p*<0.01) and 1.8-fold higher mRNA expression levels (*p*<0.01) (Figs. 1A&B) than LSECs. We also found that ethanol significantly increased CYP2E1 protein levels (1.3-fold, *p*<0.05) (Fig. 1D) in agreement with immunofluorescence images of cultured rat LSECs (Fig. 1E) and rat livers with an ethanol diet (Supplementary Fig. 1). Further, human primary LSECs showed an increase in CYP2E1 levels by ethanol treatment (2.4-fold, *p*<0.02) (Figs. 1F&G). Immunofluorescence staining of livers of endothelial-GFP reporter mice (tamoxifen inducible, Cdh5-CreERT2 mTmG mice) also showed that GFP-positive cells (liver ECs with LSECs occupying the major portion) were positive with CYP2E1 antibody, confirming CYP2E1 expression in LSECs in mouse livers as well (Fig. 1H). Collectively, these results demonstrate that LSECs express CYP2E1 and that its levels are increased by ethanol. To our knowledge, this is the first report about CYP2E1 expression in LSECs.

**Figure 1.**
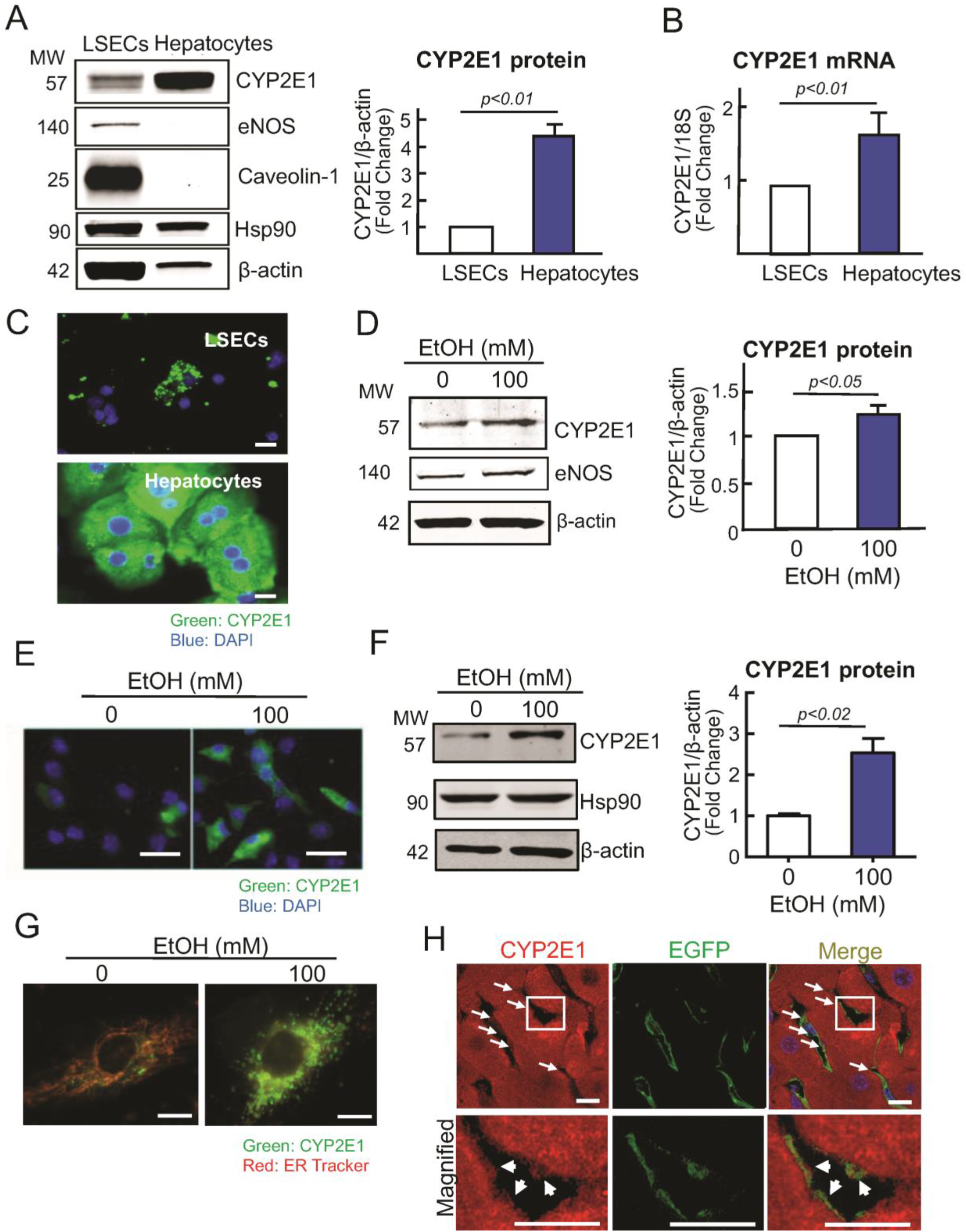
CYP2E1 is expressed in rat, mouse and human LSECs and increased by ethanol. Rat LSECs and hepatocytes were isolated and cultured for 24 hours. **(A)** Protein levels of CYP2E1, eNOS, caveolin-1, Hsp90 and β-actin were assessed by Western blotting. eNOS and caveolin-1 were used as endothelial cell markers. Hsp90 and β-actin are loading controls. n = 3. **(B)** CYP2E1 mRNA expression levels in rat LSECs and hepatocytes. n = 3. **(C)** Immunolabeling of CYP2E1 in rat LSECs and hepatocytes. Scale bar = 20 μm. **(D)** CYP2E1 protein levels in rat LSECs treated with 0 or 100 mM ethanol for 24 hours. n = 3. **(E)** Immunolabeling of CYP2E1 in rat LSECs treated with 0 or 100 mM ethanol for 24 hours. Scale bar = 50 μm. **(F)** CYP2E1 protein levels in human primary LSECs treated with 0 or 100 mM ethanol for 24 hours. n = 3. **(G)** Immunofluorescence staining of human LSECs treated with 0 or 100 mM ethanol for 24 hours. Cells were double-labeled with CYP2E1 antibody (green) and ER-Tracker (red), an endoplasmic reticulum marker. Scale bar = 20 μm. **(H)** Immunolabeling of CYP2E1 in the liver of endothelial-GFP reporter mice (tamoxifen inducible, Cdh5-CreERT2 mTmG mice). Co-localization (yellow) of CYP2E1 (red) with endothelial cells (green, GFP) was observed in liver sinusoidal areas (labelled with white arrows). Scale bar = 10 μm.

### Ethanol increases acetyl-CoA levels in LSECs

We next examined whether human LSECs could express other ethanol metabolizing enzymes. Ethanol is oxidized to acetaldehyde by cytosolic alcohol dehydrogenase 1 (ADH1) or CYP2E1. Acetaldehyde is then oxidized to acetate in the mitochondria by mitochondrial aldehyde dehydrogenase (ALDH). Finally, acetate can be converted to acetyl-CoA by acetyl-coenzyme A synthetase (ACSS) (Fig. 2A) (8). We found that human LSECs expressed ACSS1, ALDH1/2 and ADH1, but that unlike CYP2E1, their levels were not changed by ethanol (Fig. 2B). However, ethanol did increase intracellular levels of acetyl-CoA, a final product of ethanol metabolism by these enzymes (4.2-fold, *p*<0.01) (Fig. 2C). To understand the relative roles of CYP2E1 and ADH1 in ethanol metabolism in LSECs, we knocked down CYP2E1 or ADH1 in human LSECs by siRNA and treated them with ethanol. CYP2E1 and ADH1 knockdown both decreased acetyl-CoA levels, compared to a control group in response to ethanol [62% (*p*<0.005) and 40% (*p*<0.05), respectively] (Fig. 2D). Given a trend of a larger decrease in acetyl-CoA production by CYP2E1 silencing and induction of CYP2E1, but not ADH, by ethanol, these observations may suggest that CYP2E1 is more responsible for ethanol metabolism in LSECs than endogenous ADH1 under excessive alcohol intake that can lead to liver injury.

**Figure 2.**
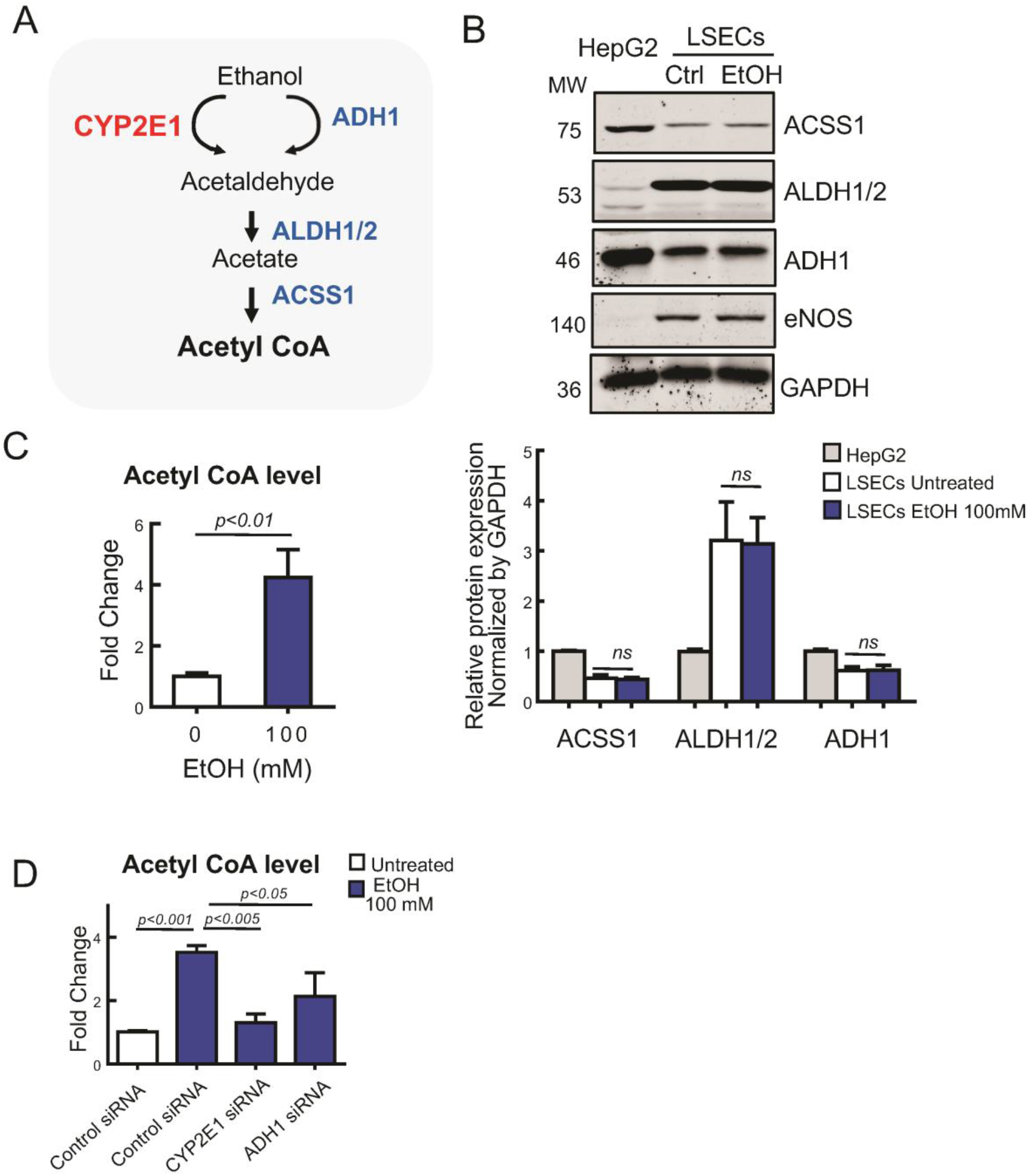
Ethanol increases acetyl-CoA levels in human primary LSECs. **(A)** A schematic diagram of ethanol metabolism. Ethanol is oxidized by cytosolic alcohol dehydrogenase 1 (ADH1) or cytochrome P450 2E1 (CYP2E1) to acetaldehyde. Acetaldehyde then enters the mitochondria to be oxidized to acetate by mitochondrial aldehyde dehydrogenase (ALDH). Acetate produced can be converted to acetyl-CoA by acetyl-coenzyme A synthetase (ACSS). **(B)** Human primary LSECs were treated with 0 or 100 mM ethanol for 24 hours. Protein levels of ACSS1, ALDH1/2, ADH1, eNOS and GAPDH were assessed in human LSECs and HepG2 by Western blotting. eNOS was used as an endothelial cell marker. GAPDH is a loading control. n = 3. **(C)** Intracellular acetyl-CoA levels in human LSECs treated with 0 or 100 mM ethanol for 24 hours. n = 4. **(D)** CYP2E1 or ADH1 expression was silenced in human LSECs by siRNA. Twenty-four hours after transfection, human LSECs were treated with 0 or 100 mM ethanol for 24 hours. Intracellular acetyl-CoA levels were then assessed. n = 3.

### Ethanol decreases NO production and increases Hsp90 acetylation in the presence of CYP2E1

LSECs produce NO by the action of eNOS to regulate intrahepatic vascular tone in normal conditions. Decreased production of NO is a characteristic of LSEC dysfunction (20). Ethanol significantly decreased NO production in human LSECs (39%, *p*<0.01) (Fig. 3A) without changing eNOS protein levels (Fig. 3B). A similar result was observed in bovine aortic endothelial cells (BAECs), frequently used ECs for the study of eNOS (Supplementary Figs. 2A&B). These observations indicate that the decreased production of NO by ethanol is due to compromised eNOS activity.

**Figure 3.**
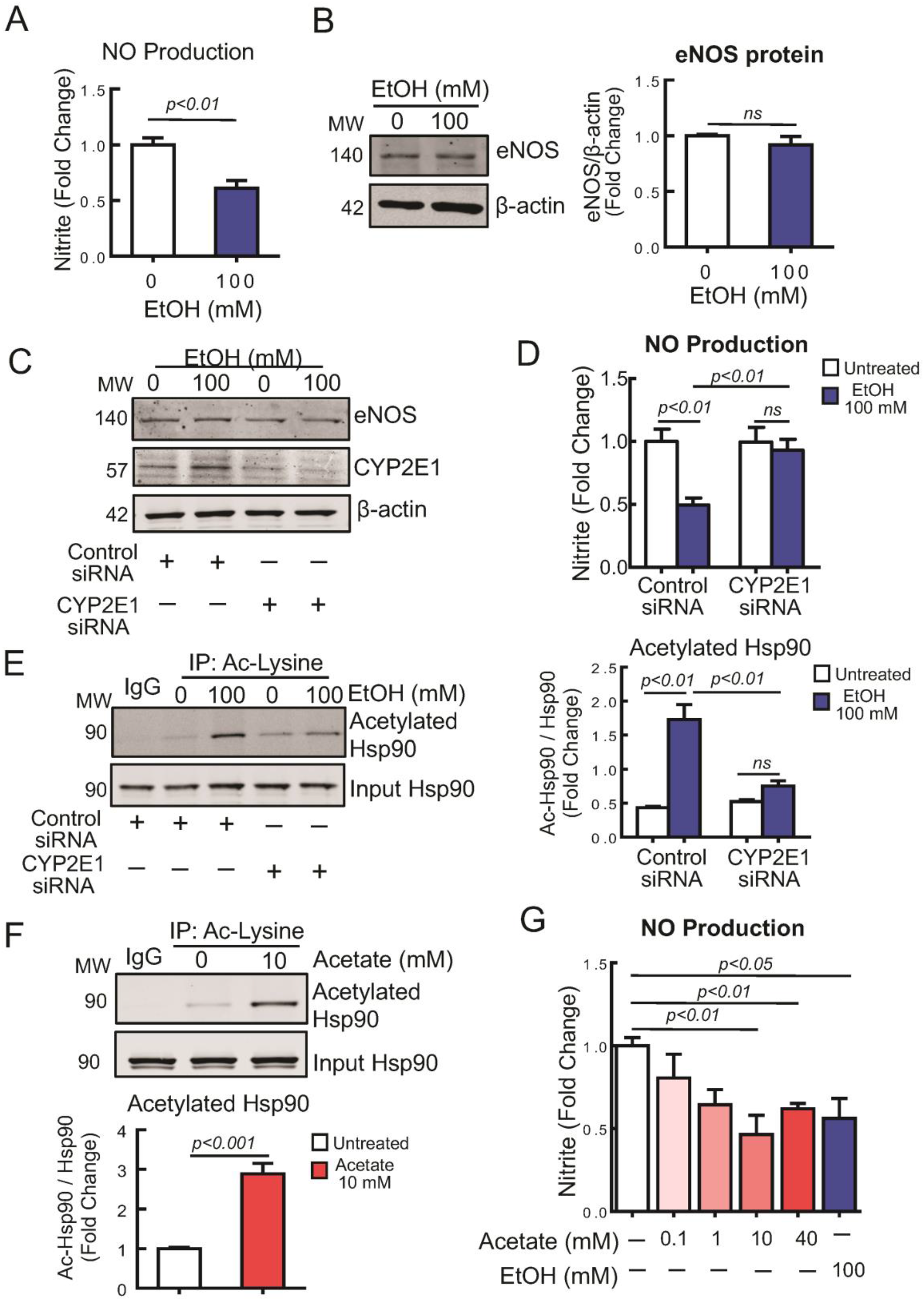
Ethanol decreases NO production and increases Hsp90 acetylation in the presence of CYP2E1. **(A)** Nitrite (an end product of NO) levels in culture medium of human LSECs treated with 0 or 100 mM ethanol for 24 hours. n = 6. **(B)** eNOS protein levels in human LSECs treated with 0 or 100 mM ethanol. β-actin is a loading control. n = 3. **(C&D)** CYP2E1 expression was silenced in human LSECs by siRNA. Twenty-four hours after transfection, human LSECs were treated with 0 or 100 mM ethanol for 24 hours. Protein levels of eNOS and CYP2E1 were assessed by Western blotting. n = 3. Nitrite levels in culture medium were determined. n = 6. **(E)** Cell lysates of human LSECs were used for immunoprecipitation (IP) with acetyl-lysine antibody to examine Hsp90 acetylation in response to ethanol. Western blotting was performed to detect acetylated Hsp90 in IP samples. Input Hsp90 is total Hsp90 in input samples used for this IP experiment. n = 4. **(F)** Human LSECs were treated with 0 or 10 mM acetate for 24 hours. Cell lysates were used for IP with acetyl-lysine antibody, followed by Western blotting to detect acetylated Hsp90 in IP samples and total Hsp90 in input samples. n = 5. **(G)** Human LSECs were treated with ethanol or acetate with indicated concentrations for 24 hours. Nitrite levels in culture medium were determined. n = 4.

To understand the role of CYP2E1 in LSEC dysfunction in response to ethanol, we knocked down CYP2E1 in human LSECs by siRNA and treated them with ethanol. Suppression of CYP2E1 did not change eNOS protein levels (Fig. 3C) but recovered NO production (Fig. 3D). Since CYP2E1 metabolizes ethanol and leads to the production of acetyl-CoA, a substrate for protein acetylation, we hypothesized that CYP2E1 could contribute to protein acetylation through acetyl-CoA production. As shown in Fig. 3E, ethanol significantly increased Hsp90 acetylation (4-fold, *p*<0.01) in human LSECs, which was reversed by CYP2E1 knockdown. Similarly, ethanol significantly increased Hsp90 acetylation (1.54-fold, *p*<0.05) in BAECs (Supplementary Fig. 2C). We then used acetate to recapitulate the effect of acetyl-CoA on Hsp90 acetylation and NO production in human LSECs. Similar to ethanol, acetate increased Hsp90 acetylation (Fig. 3F) (2.9-fold, *p*<0.01) and reduced NO production in a dose-dependent manner (Fig. 3F). Collectively, these results indicate that ethanol decreases NO production and increases Hsp90 acetylation in the presence of CYP2E1.

### Ethanol-driven Hsp90 acetylation decreases its interaction with eNOS

Hsp90 is a positive regulator of eNOS activity (14). However, whether Hsp90 acetylation influences eNOS activity is not known. We hypothesized that Hsp90 acetylation could decrease its interaction with eNOS and reduce NO production (Fig. 4A). As shown in Fig. 4B, ethanol decreased Hsp90’s interaction with eNOS in human LSECs (25%, *p*<0.02), but this interaction was restored by CYP2E1 knockdown, indicating that decreased interaction of Hsp90 with eNOS by ethanol is CYP2E1-dependent. A similar pattern was observed in BAECs with both ethanol and acetaldehyde decreasing Hsp90’s interaction with eNOS and reducing NO production (Supplementary Figs. 3A&B).

**Figure 4.**
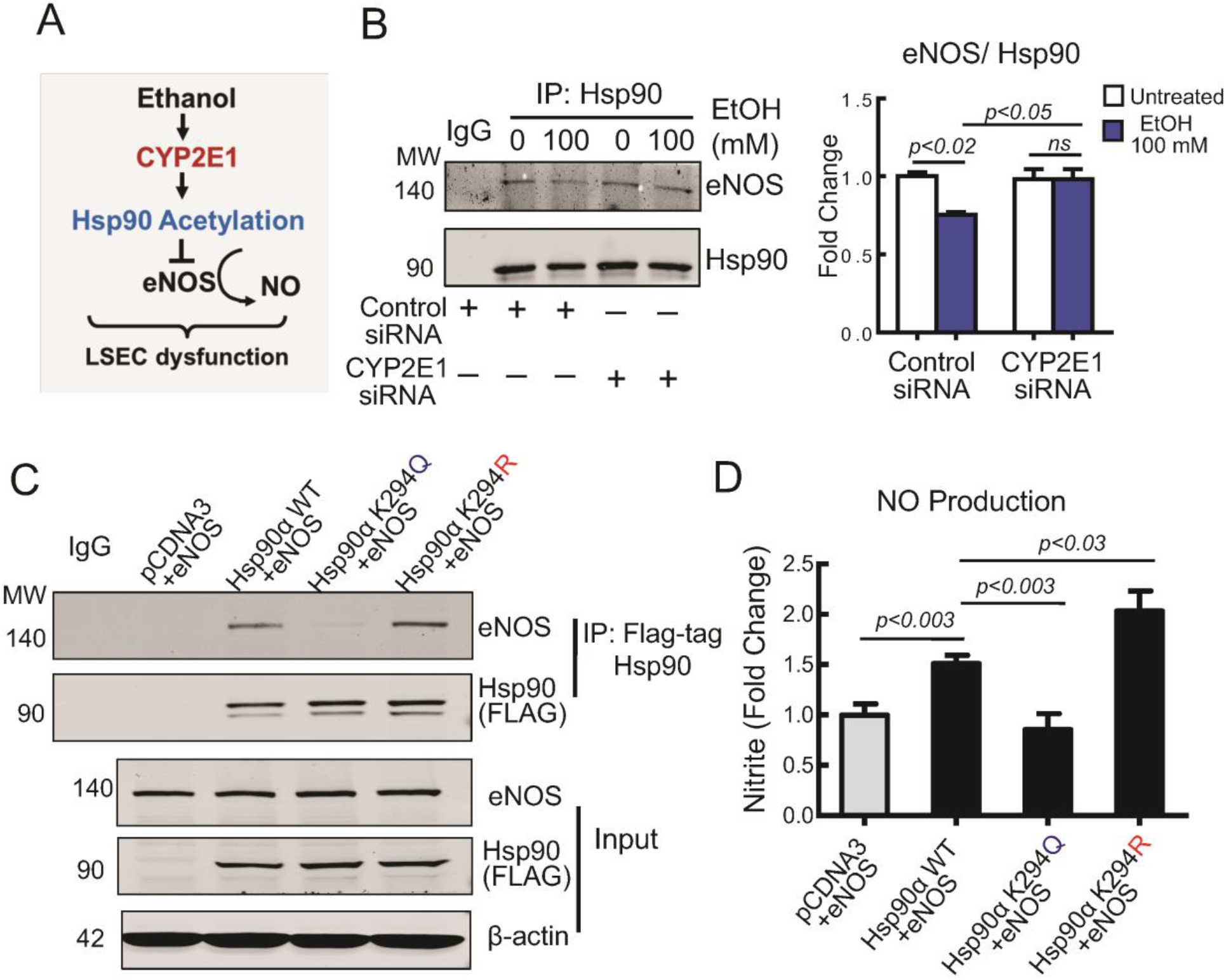
Ethanol-driven Hsp90 acetylation decreases its interaction with eNOS. **(A)** Potential mechanism of ethanol-induced LSEC dysfunction. Ethanol increases CYP2E1 expression in LSECs and promotes Hsp90 acetylation, which interferes Hsp90’s interaction with eNOS and decreases eNOS-derived NO production. **(B)** Human LSECs were treated with CYP2E1 siRNA for 24 hours and then with 0 or 100 mM ethanol for another 24 hours. Cell lysates were used for immunoprecipitation (IP) with mouse IgG (control) or Hsp90 antibody. Western blotting was performed to detect Hsp90 and eNOS. n = 3. **(C&D)** COS7 cells were transfected with eNOS and pCDNA3 vector (control), WT Hsp90α or Hsp90α mutants (K294Q and K294R). A mutation to glutamine (Q) represents a hyper-acetylated state of Hsp90, while an arginine (R) mutant cannot be acetylated. Twenty-four hours after transfection, COS7 cells were further cultured for 24 hours, and cell lysates and medium were collected. **(C)** Cell lysates were immunoprecipitated (IP) with FLAG-tag antibody to pull-down Hsp90 to detect eNOS that interacted with Hsp90. n = 3. **(D)** Nitrite (an end product of NO) levels in culture medium were determined. n = 8.

We further confirmed the role of Hsp90 acetylation in its interaction with eNOS and NO production by mutating its specific lysine residue for acetylation. A mutation to glutamine (Q) represents a hyper-acetylated state of Hsp90, while an arginine (R) mutant cannot be acetylated. We assessed Hsp90’s interaction with eNOS and NO production in COS7 cells, which express Hsp90, but not eNOS (14, 21, 22). Thus, eNOS was transfected along with the Hsp90 mutants. As shown in Fig. 4C, the non-acetylation mutant of Hsp90α (K294R) increased its interaction with eNOS by 2-fold compared with WT, or even greater compared with the hyper-acetylation mutant (K294Q). Further, the non-acetylation mutant (K294R) produced NO more than any other type, whereas the hyper-acetylation mutant (K294Q) did not increase NO production (Fig. 4D). These observations strongly support our hypothesis that the state of Hsp90 acetylation influences its interaction with eNOS and NO production.

### HDAC6 inhibition increases Hsp90 acetylation and decreases NO production

HDAC6 is a major enzyme to regulate Hsp90 acetylation. HDAC6 decreases Hsp90 acetylation levels by removing an acetyl group (23). We used an HDAC6 inhibitor, trichostatin A (TSA), to examine its effect on Hsp90 acetylation (Fig. 5A). As expected, HDAC6 inhibition by TSA significantly increased Hsp90 acetylation (2.5-fold, *p*<0.02) (Fig. 5B). HDAC6 inhibition also decreased Hsp90’s interaction with eNOS (70%, *p*<0.05) (Fig. 5C) and NO production (25%, *p*<0.001) (Fig. 5D). Since TSA did not change eNOS protein levels (Fig. 5E), the decreased production of NO may be attributable to the change in the interaction between Hsp90 and eNOS that influences eNOS activity. Collectively, Hsp90 acetylation promoted by HDAC6 inhibition decreases Hsp90’s interaction with eNOS and inhibits NO production. Conversely, HDAC6 overexpression could reduce Hsp90 acetylation and facilitate Hsp90’s interaction with eNOS in LSECs, thereby increasing NO production in LSECs and protecting LSECs from ethanol-induced dysfunction.

**Figure 5.**
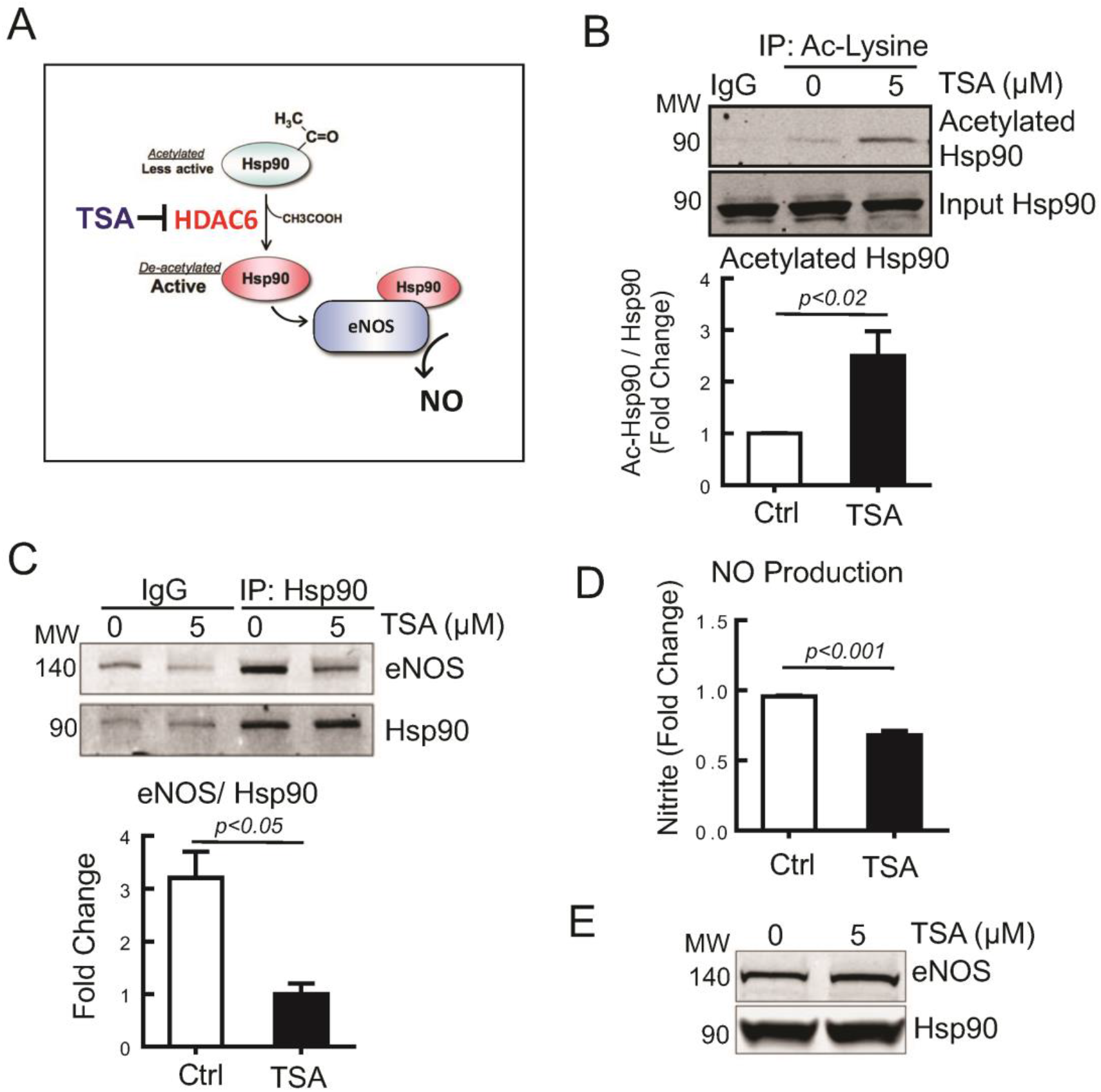
HDAC6 inhibition increases Hsp90 acetylation and decreases NO production in BAECs. **(A)** The mechanism of Hsp90 acetylation by trichostatin A (TSA), a histone deacetylase 6 (HDAC6) inhibitor. The level of Hsp90 acetylation is regulated by HDAC6, which removes an acetyl group from acetylated Hsp90. TSA can increase Hsp90 acetylation by inhibiting HDAC6. **(B - E)** Bovine aortic endothelial cells (BAECs), commonly used ECs for the study of eNOS, were treated with TSA for 24 hours. **(B)** Cell lysates of BAECs were used for immunoprecipitation (IP) with acetyl-lysine antibody, followed by Western blotting to detect acetylated Hsp90 in IP samples and total Hsp90 in input samples. n = 4. **(C)** Cell lysates were used for IP with Hsp90 antibody, followed by Western blotting to detect Hsp90 and eNOS. n = 3. **(D)** Nitrite (an end product of NO) levels in culture medium were determined. n = 7. **(E)** Protein levels of eNOS and Hsp90 were assessed by Western blotting. n = 3.

### HDAC6 overexpression in liver ECs protects Hsp90’s interaction with eNOS against ethanol-induced acetylation *in vivo*

We hypothesized that inhibiting Hsp90 acetylation by overexpressing HDAC6 in liver ECs could restore EC function in alcoholic liver disease (Fig. 6A). To this end, we generated adeno-associated virus 8 (AAV8)-HDAC6 (AAV8-EFS-DIO-mHDAC6), which allows HDAC6 expression only in the presence of Cre-recombinase. AAV-HDAC6 or AAV-control (Ctrl) (AAV8-EF1a-DIO-mKate2) was injected to tamoxifen-induced Cdh5-CreERT2 mice (EC-specific Cre mice) 3 weeks before feeding an ethanol diet for 10 days. A control group received AAV injection similarly but was kept on a control diet (Fig. 6B). Because AAV8 has a significantly greater liver targeting ability (24), this approach permits liver EC-specific HDAC6 overexpression without influencing ECs in other organs. We confirmed HDAC6 overexpression in liver ECs, particularly LSECs, as indicated by co-localization of HDAC6 expression with PECAM-1, an EC marker, in mice administered AAV-HDAC6, compared to mice given AAV-Ctrl (Fig. 6C). We also confirmed that both HDAC6 mRNA and protein levels were elevated in AAV-HDAC6 mouse livers (Figs. 6D&E). Ethanol feeding did not change HDAC6 protein levels, but significantly increased CYP2E1 (Fig. 6E). As shown in Fig. 6F, ethanol feeding significantly increased Hsp90 acetylation, but HDAC6 overexpression by AAV-HDAC6 had it reduced We also observed that an interaction between Hsp90 and eNOS was diminished by ethanol feeding, but was recovered by HDAC6 overexpression (Fig. 6G). These results demonstrate that HDAC6 overexpression in liver ECs can reduce ethanol-driven Hsp90 acetylation and increase the Hsp90-eNOS interaction in ethanol fed mice.

**Figure 6.**
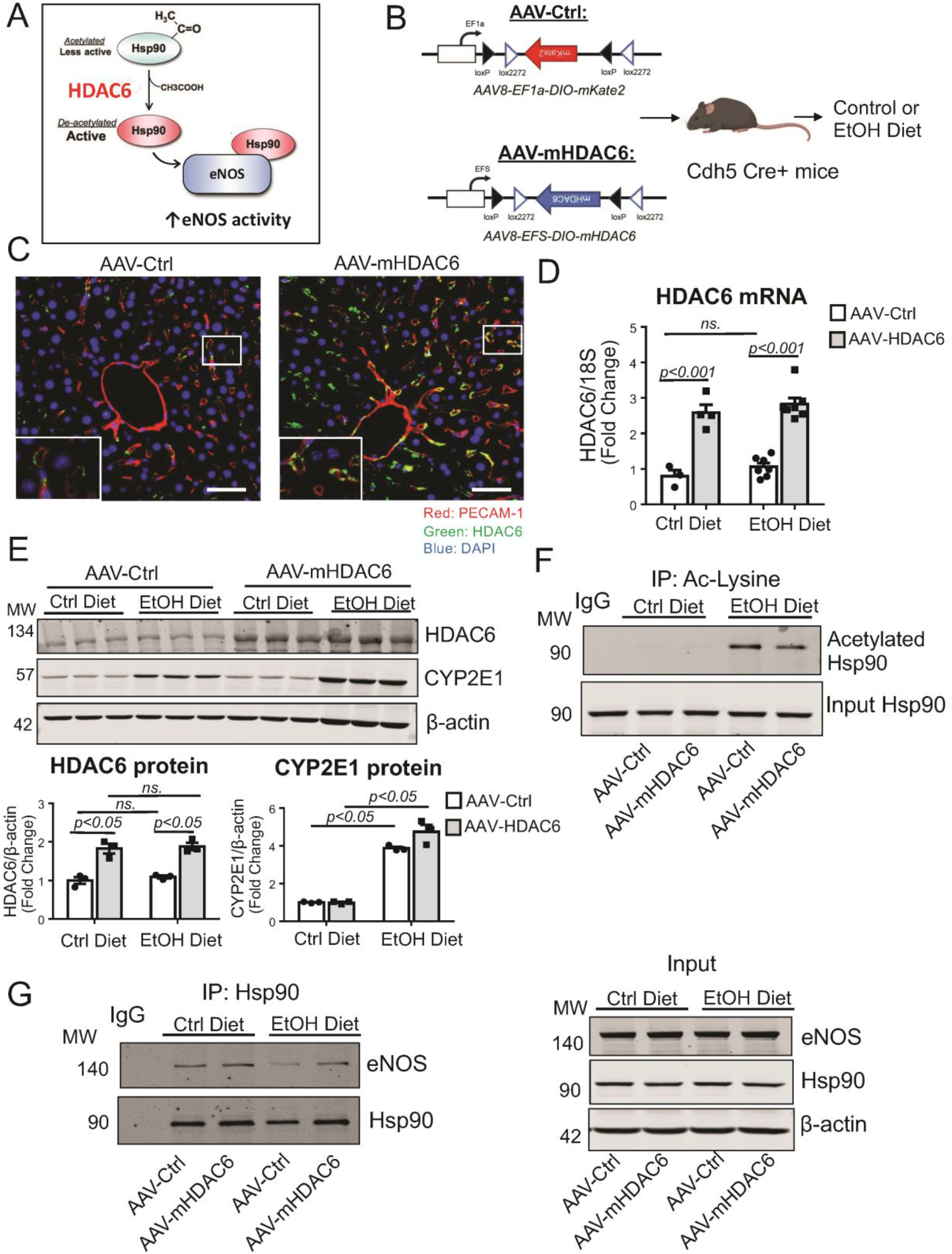
*In vivo* HDAC6 overexpression in liver ECs recovers Hsp90’s interaction with eNOS in ethanol fed mice. **(A)** Potential mechanism of Hsp90-eNOS interaction regulated by histone deacetylase 6 (HDAC6). HDAC6 overexpression in liver ECs may decrease Hsp90 acetylation and enhance Hsp90-eNOS interaction in alcoholic liver disease. **(B)** An *in vivo* experiment workflow and construct maps of adeno-associated virus (AAV)-Ctrl (AAV8-EF1a-DIO-mKate2) and AAV-HDAC6 (AAV8-EFS-DIO-mHDAC6). Tamoxifen induced Cdh5-CreERT2 mice (EC-specific Cre-recombinase expression) were injected with AAV-Ctrl or AAV-HDAC6 through the retro-orbital sinus. Three weeks later, these mice were fed with an ethanol diet for 10 days, followed by an oral ethanol gavage. Nine hours after gavage, samples were collected. A control group was injected with AAV constructs similarly. Three weeks later, they were fed with a control diet for 10 days and sacrificed for sample collection. **(C)** Immunolabeling of HDAC6 (green) and PECAM-1 (red, an EC marker) in liver specimens from Cdh5-CreERT2 mice injected with AAV-Ctrl or AAV-HDAC6 and fed with a control diet. Scale bar = 50 μm. **(D)** HDAC6 mRNA expression levels in mouse livers. n = 3 in AAV-Ctrl group and n = 4 in AAV-HDAC6 group for Control diet, n = 7 per group for Ethanol diet. **(E)** Protein levels of HDAC6 and CYP2E1 were assessed by Western blotting. β-actin is a loading control. n = 3. **(F)** Liver tissue lysates were immunoprecipitated with acetyl-lysine antibody, followed by Western blotting to detect total and acetylated Hsp90. **(G)** Liver tissue lysates were immunoprecipitated with eNOS antibody, followed by Western blotting to detect eNOS and Hsp90 (left panel). Protein levels of eNOS and Hsp90 in input samples were assessed by Western blotting (right panel).

### HDAC6 overexpression in liver ECs reduces alcohol-induced liver injury

We also investigated the effect of HDAC6 overexpression in liver ECs on alcohol-induced liver injury. Plasma ALT levels were significantly elevated by ethanol feeding but decreased by HDAC6 overexpression (Fig. 7A). Similarly, neutrophil infiltration assessed by qPCR and immunolabeling of myeloperoxidase (MPO) was significantly decreased in AAV-HDAC6 mice compared to AAV-Ctrl mice (Figs. 7B&C). Histologically, hepatic inflammation was increased by ethanol feeding but alleviated by HDAC6 overexpression (Fig. 7D). Immunolabeling of CK8 also showed reduced hepatic injury by HDAC6 overexpression (Fig. 7E). Further, the levels of pro-inflammatory cytokines including tumor necrosis factor (TNF)-α, interleukin (IL)-1β and IL-6 were significantly lower in AAV-HDAC6 mice than AAV-Ctrl mice (Fig. 7F, Supplementary Fig. 4). All these results indicate that HDAC6 overexpression in liver ECs protects mice from alcohol-induced liver injury.

**Figure 7.**
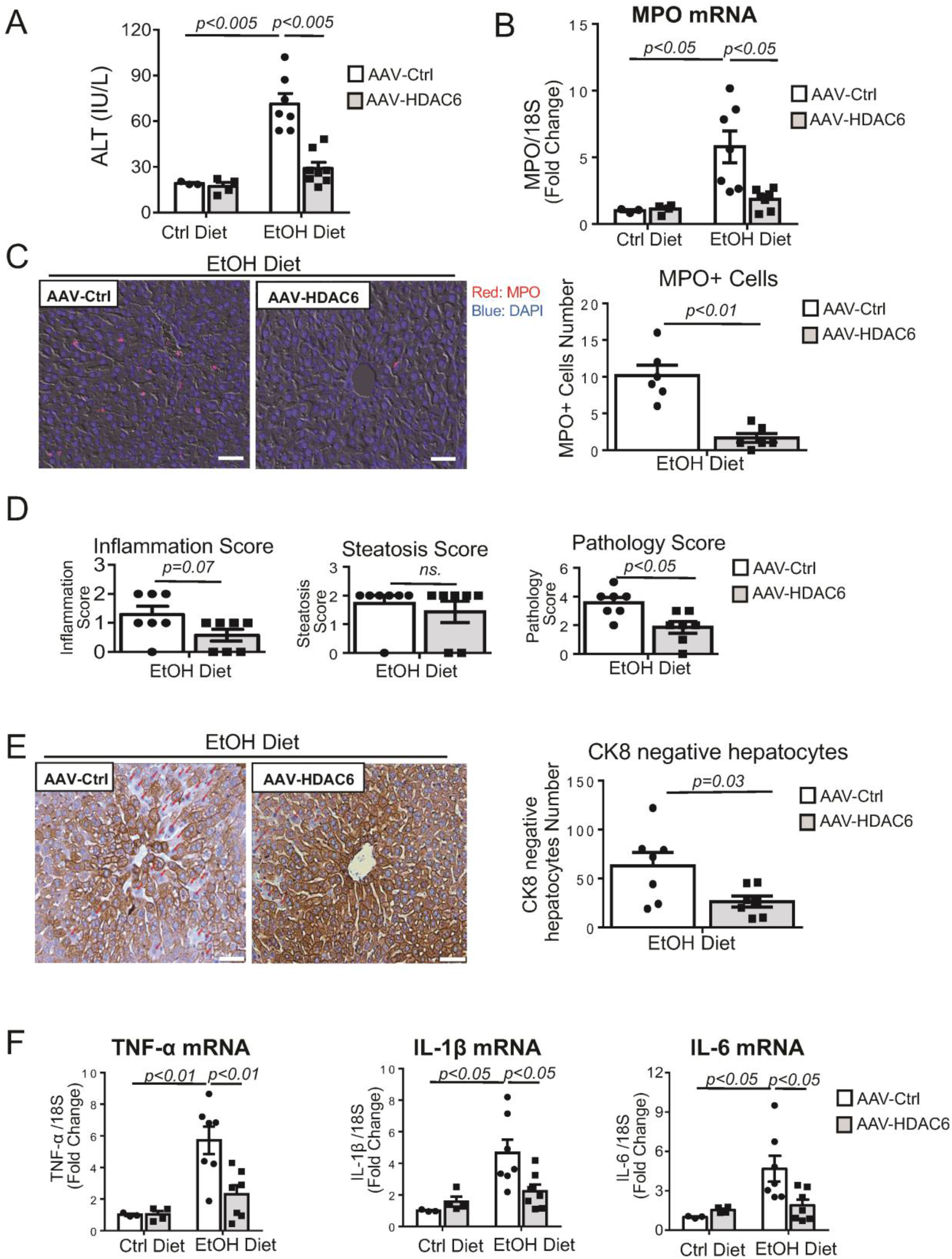
*In vivo* HDAC6 overexpression in liver ECs reduces alcohol-induced liver injury. Cdh5-CreERT2 mice were injected with AAV-Ctrl or AAV-HDAC6. Three weeks later, these mice were fed with an ethanol diet for 10 days, followed by an oral ethanol gavage. Nine hours after gavage, samples were collected. A control group received AAV injection similarly and had a control diet for 10 days, followed by sample collection. **(A)** Serum ALT levels. n = 3 in AAV-Ctrl group and n = 4 in AAV-HDAC6 group for Control diet, n = 7 per group for Ethanol diet. **(B)** Myeloperoxidase (MPO) mRNA expression levels in mouse livers. n = 3 in AAV-Ctrl group and n = 4 in AAV-HDAC6 group for Control diet, n = 7 per group for Ethanol diet. **(C)** Immunolabeling of MPO in mouse livers. Scale bar = 50 μm. n = 6. **(D)** Histological scores. Scale bar = 50 μm. n = 7. **(E)** CK8 staining of mouse livers. CK8-negative hepatocytes were marked by red arrows and quantified. Scale bar = 50 μm. n = 7. **(F)** mRNA expression levels of TNF-α, IL-1β and IL-6 in mouse livers. n = 3 in AAV-Ctrl group and n = 4 in AAV-HDAC6 group for Control diet, n = 7 per group for Ethanol diet.

## Discussion

Alcohol metabolism has not been described in LSECs. In this study, we have revealed CYP2E1 and ADH1, key enzymes in alcohol metabolism, are constitutively expressed in LSECs. We have also shown that ethanol increases CYP2E1 levels (but not ADH1 levels) as well as intracellular levels of acetyl-CoA, a final product of alcohol metabolism, in LSECs. To our knowledge, this is the first study to demonstrate CYP2E1/ADH1 expression and alcohol metabolism in LSECs.

Several studies have reported LSEC dysfunction in relation to alcoholic liver disease (4, 5, 25) with some of them indicating that decreased hepatic NO production under excessive alcohol intake is associated with alcohol-induced liver injury (26, 27). However, the mechanism by which alcohol consumption decreases NO production remains unknown. In this study, we have determined diminished interaction of Hsp90 with eNOS as the underlying mechanism of decreased NO production and liver injury under excessive alcohol consumption.

Our study has shown that ethanol increases Hsp90 acetylation and inhibits its interaction with eNOS. Because eNOS levels were not changed by ethanol, decreased eNOS activity caused by diminished interaction with Hsp90 due to its acetylation is responsible for decreased NO production in LSECs. Our experiment using Hsp90 constructs with a mutation in the acetylation site confirmed the effect of Hsp90 acetylation on its interaction with eNOS and eNOS-derived NO production. This is a new mechanism of eNOS regulation by Hsp90 acetylation. More importantly in our context, Hsp90 acetylation occurs in the presence of CYP2E1 or acetate, highlighting the importance of alcohol metabolism in LSECs.

In normal conditions, ADH1 enzyme, by metabolizing alcohol, may also facilitate Hsp90 acetylation to some degree without causing LSEC dysfunction. Under excess alcohol intake, since ADH1 is not induced by ethanol in LSECs, significantly upregulated CYP2E1 increases Hsp90 acetylation and impairs eNOS-derived NO production in LSECs.

Our proof of concept experiments manipulating HDAC levels have also confirmed the relationship between Hsp90 acetylation and its interaction with eNOS. *In vitro*, HDAC6 inhibition by trichostatin A (TSA) significantly increased Hsp90 acetylation, decreased its interaction with eNOS and decreased NO production without any changes in eNOS protein levels. *In vivo*, overexpression of HDAC6 in liver ECs by AAV8-mediated gene delivery decreased Hsp90 acetylation and increased its interaction with eNOS in mouse livers. Most importantly, overexpression of HDAC6 reduced alcohol-induced liver injury, as indicated by decreased plasma ALT levels, myeloperoxidase expression and liver inflammation. Expression of many pro-inflammatory cytokines including TNF-α, IL-1β and IL-6 were also significantly reduced. Since LSECs account for approximately 90% of liver ECs, it is reasonable to think that the protective effect of HDAC6 overexpression can be mediated by LSECs more than any other liver endothelial cells. In addition, AAV8 is a serotype that is preferentially expressed in the liver, which could avoid a confounding effect of HDAC6 on endothelial cells in other organs. Therefore, these results strongly suggest that restoring LSEC function is an important therapeutic strategy for alcoholic liver disease.

The role of HDAC6 in LSECs and in response to ethanol remains to be elucidated (28). In WIF-B cells, ethanol treatment slightly decreased HDAC6 protein levels (28). In a recent study, however, binge drinking did not change gene expression levels of HDAC6 in rat livers (29). We did not observe any changes in either HDAC6 protein or gene expression levels in ethanol fed mouse livers. Because these *in vivo* studies including ours determined HDAC6 levels in whole liver lysates, it is still possible that alcohol may negatively regulate HDAC6 levels in LSECs.

It should also be mentioned that some studies have regarded HDAC6 as pro-oncogenic and investigated an HDAC6 inhibitor for the treatment of hepatocellular carcinoma (HCC) (30, 31). For example, a recent study (31) has described that deletion of HDAC6 in T helper 17 (Th17) cells impedes HCC growth by enhancing the production of antitumor cytokines and CD8 T cell mediated antitumor response in mice. However, other studies have shown that HDAC6 is a tumor suppressor for HCC (32–34). It is reported that low expression of HDAC6 in liver cancer specimens is highly associated with poor prognosis of HCC patients (32). Sustained HDAC6 overexpression is shown to trigger autophagic death of cancer cells and reduce the *in vivo* tumor growth rate in a mouse xenograft model. Another study (33) has demonstrated that HDAC6 is a tumor suppressor for HCC by inhibiting canonical Wnt/b-catenin signaling, which is known to be involved in hepatocarcinogenesis and metastasis. Our study has shown that liver EC-specific HDAC6 overexpression is beneficial for alcohol-induced liver injury.

Protein acetylation has been described in relation to ethanol exposure. It is reported that ethanol exposure induces global protein hyperacetylation, contributing to alcohol-induced hepatotoxicity (28, 35, 36). Chronic ethanol exposure promotes α-tubulin acetylation, which enhances microtubule stability, impairs lipid droplet motility and thus causes functional defects of hepatocytes (37). Histone acetylation has also been observed in isolated hepatocytes and rat livers in response to both acute and chronic ethanol exposure, which could lead to epigenetic alterations in hepatocytes (38, 39). Therefore, besides deacetylation of Hsp90, it is possible that HDAC6 might have deacetylated other proteins in liver ECs and helped to ameliorate alcohol-induced liver injury. With acetyl-CoA as a product of alcohol metabolism, protein acetylation in alcoholic liver disease may be an interesting area of investigation, which would advance our understanding of molecular mechanisms leading to liver injury.

Several studies have addressed eNOS acetylation with inconsistent results regarding its effect. For example, aspirin and acetylating analogs were reported to activate eNOS by directly acetylating eNOS (40). Sirtuin1 (SIRT1) enhanced eNOS activity by directly deacetylating eNOS at lysines 496 and 506 (40–42), while histone deacetylase 3 (HDAC3) deacetylated eNOS and decreased NO production by reducing an association of eNOS with calmodulin, a positive regulator of eNOS activity (43). In our study, ethanol did not change eNOS acetylation (data not shown), reinforcing that diminished eNOS activity is attributable to Hsp90 acetylation by CYP2E1-mediated alcohol metabolism.

Although CYP2E1 expression in LSECs had not been known, some other endothelial cells, such as human umbilical endothelial cells (HUVECs) and brain microvascular endothelial cells, were reported to express CYP2E1 (44, 45). The functional significance of CYP2E1 in HUVECs is not known (44). In brain microvascular endothelial cells, ethanol induces CYP2E1 expression and increases oxidative stress, resulting in blood-brain barrier dysfunction (45). CYP2E1 generates reactive oxygen species (ROS) as a byproduct of alcohol metabolism, which can cause tissue injury (8, 46). Deletion of CYP2E1 in mice showed reduced alcohol-induced liver injury (47), while transgenic overexpression of CYP2E1 exacerbated alcohol-induced liver injury in mice (48). Thus, increased ROS as a result of CYP2E1 induction could also contribute to LSEC dysfunction. Further, ROS, such as superoxide radicals (O2-), can spontaneously react with NO to form a harmful peroxynitrite, further promoting cellular damage. This could also lead to diminished bioavailability of NO, contributing to decreased NO levels in LSECs in another way.

Of note, it is reported that production of extracellular vesicles (EVs) is significantly increased in patients with alcoholism and rats with ethanol exposure and that these EVs are highly enriched with CYP2E1 (49). Given the close proximity between LSECs and hepatocytes in the hepatic microenvironment, it is highly possible that hepatocyte-derived CYP2E1 proteins could be transported to LSECs via EVs and contribute to LSEC dysfunction. Therefore, deletion of the CYP2E1 gene in LSECs may not be a useful measure for LSEC protection against ethanol exposure. Rather, targeting downstream effectors of CYP2E1, such as Hsp90 acetylation, would be a more plausible therapeutic approach.

In conclusion, alcohol metabolism by CYP2E1 in LSECs contributes to liver injury. Restoration of LSEC function, particularly maintaining NO levels in LSECs, may be important for ameliorating alcohol-induced liver injury. In this regard, HDAC6 overexpression specifically in LSECs with AAV8-mediated gene delivery may represent a new therapeutic strategy.

## Acknowledgements

We would like to thank Dr. Samuel Varman (Yale University) for providing us with liver samples from ethanol fed rats. Drs. Len Neckers and Brad Scroggins (National Cancer Institute) for Hsp90α acetylation mutants and technical support. We would also like to thank Kathy Harry at the Yale Liver Center Cell Isolation Core Facility.

## Supplemental Materials

### Supplementary Figures

**Supplementary Figure 1.**
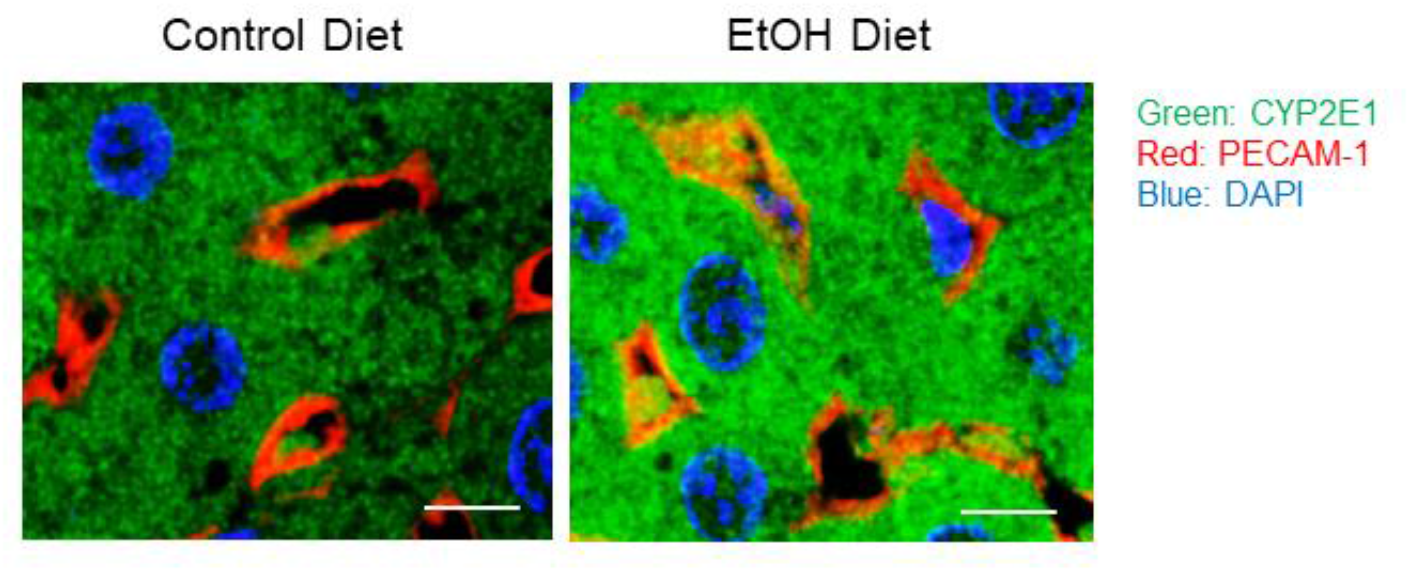
Ethanol increases CYP2E1 expression in LSECs in the liver. Immunofluorescence images of rat livers with control or ethanol diet (NIAAA model). An increase in CYP2E1 in LSECs was indicated by increased yellow color [an overlap of CYP2E1 (green) and PECAM1 (red, an EC marker)] in the image from ethanol fed rats. Scale bar = 10 μm.

**Supplementary Figure 2.**
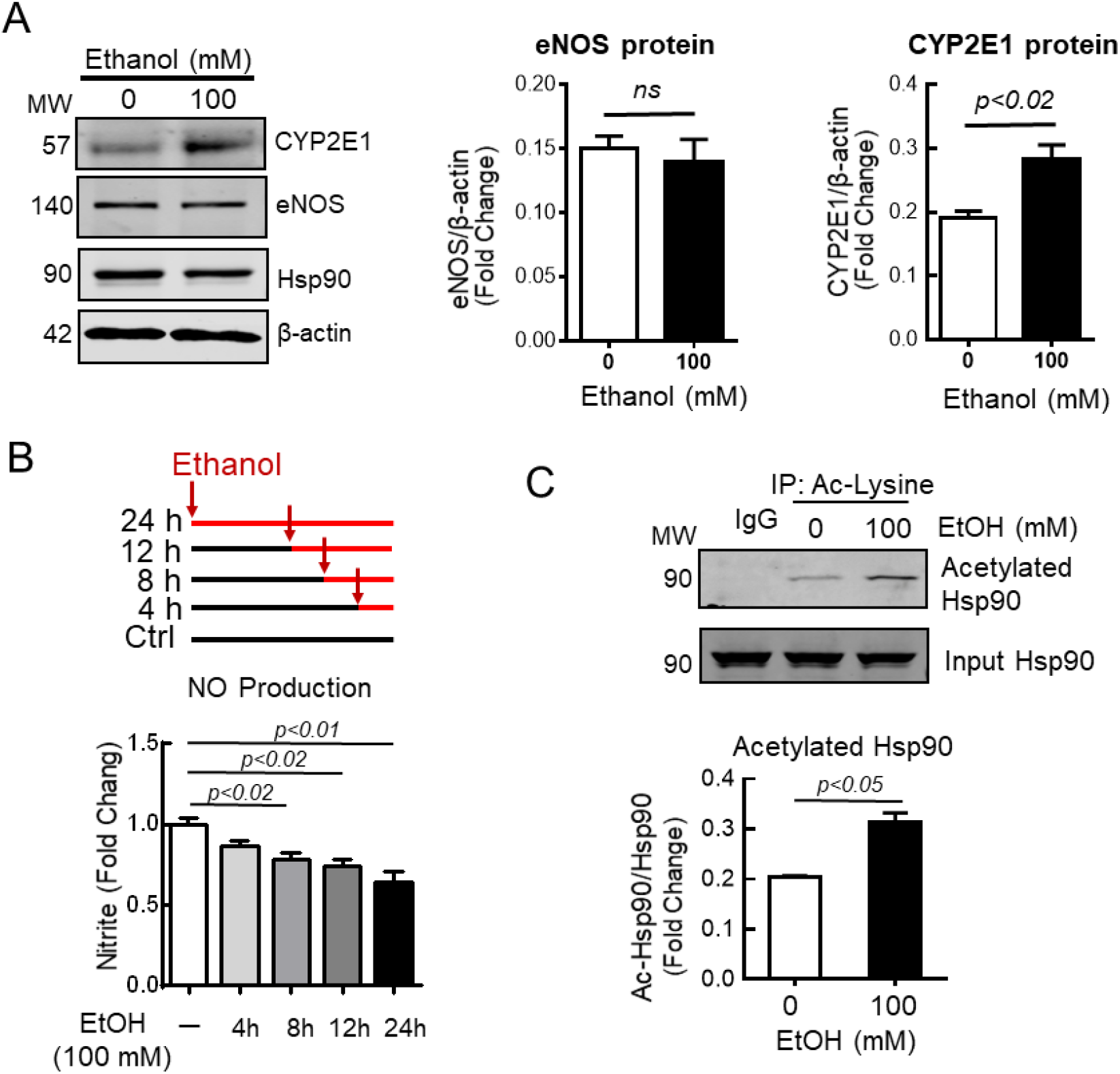
Ethanol decreases NO production and increases Hsp90 acetylation in BAECs. **(A)** Bovine aortic endothelial cells (BAECs) were treated with 0 or 100 mM ethanol for 24 hours. Protein levels of CYP2E1 and eNOS were assessed by Western blotting. Hsp90 and β-actin are loading controls. n = 3. **(B)** 24 hour-accumulations of nitrite (an end product of NO) in culture medium of BAECs treated with 100 mM ethanol for different durations (0, 4, 8, 12 and 24 hours) were determined. n = 6. **(C)** BAECs were treated with 0 or 100 mM ethanol for 24 hours. Cell lysates were used for immunoprecipitation with acetyl-lysine antibody, followed by Western blotting to detect total and acetylated Hsp90. n = 3.

**Supplementary Figure 3.**
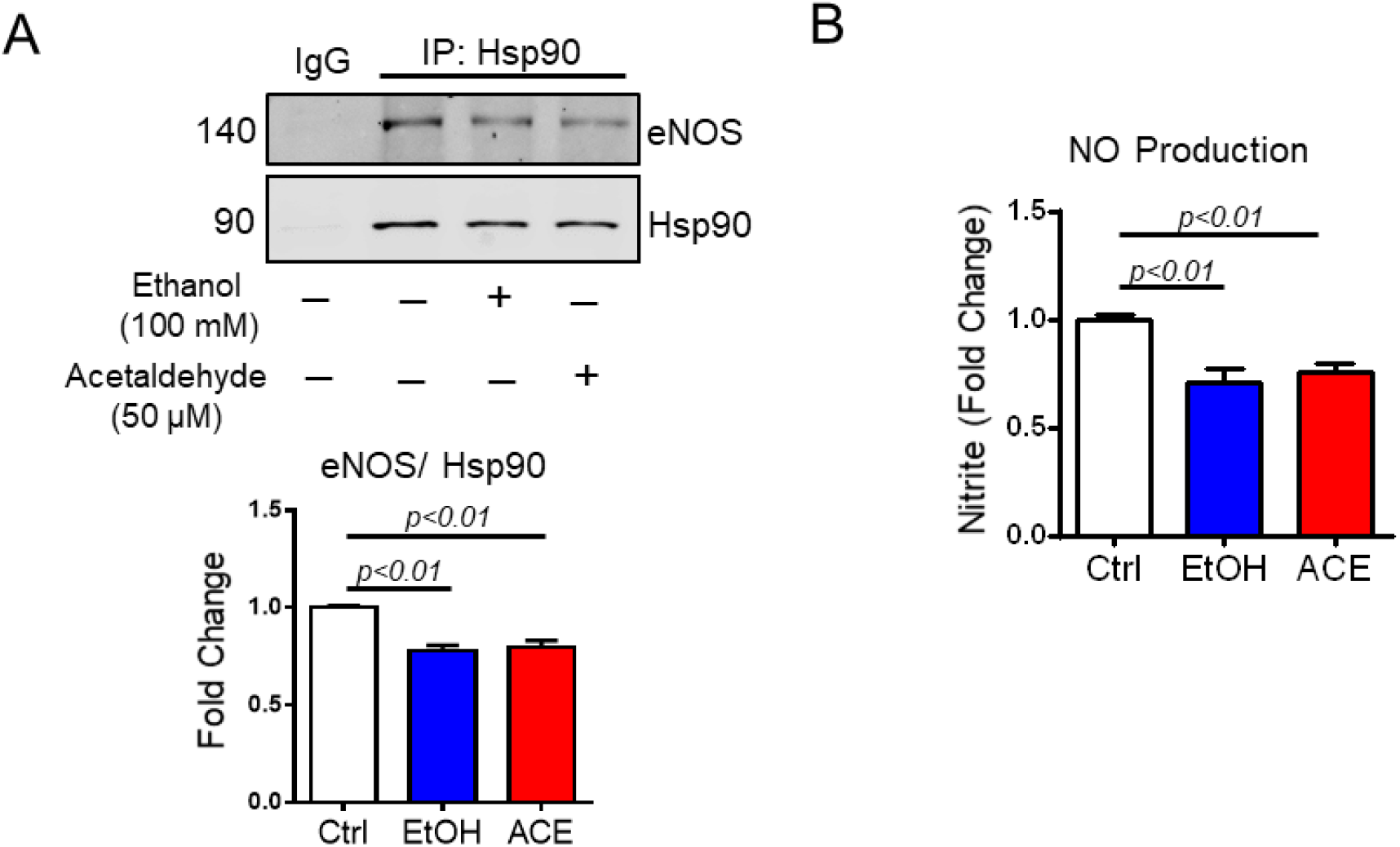
Ethanol and acetaldehyde decrease Hsp90’s interaction with eNOS in BAECs. **(A)** Bovine aortic endothelial cells (BAECs) were treated with 50 μM acetaldehyde (ACE) or 100 mM ethanol for 24 hours. Cell lysates were used for immunoprecipitation with Hsp90 antibody, followed by Western blotting to detect Hsp90 and eNOS. n = 3. **(B)** BAECs were treated with 50 μM acetaldehyde (ACE) or 100mM ethanol for 24 hours. Nitrite (an end product of NO) levels in culture medium were determined. n = 6.

**Supplementary Figure 4.**
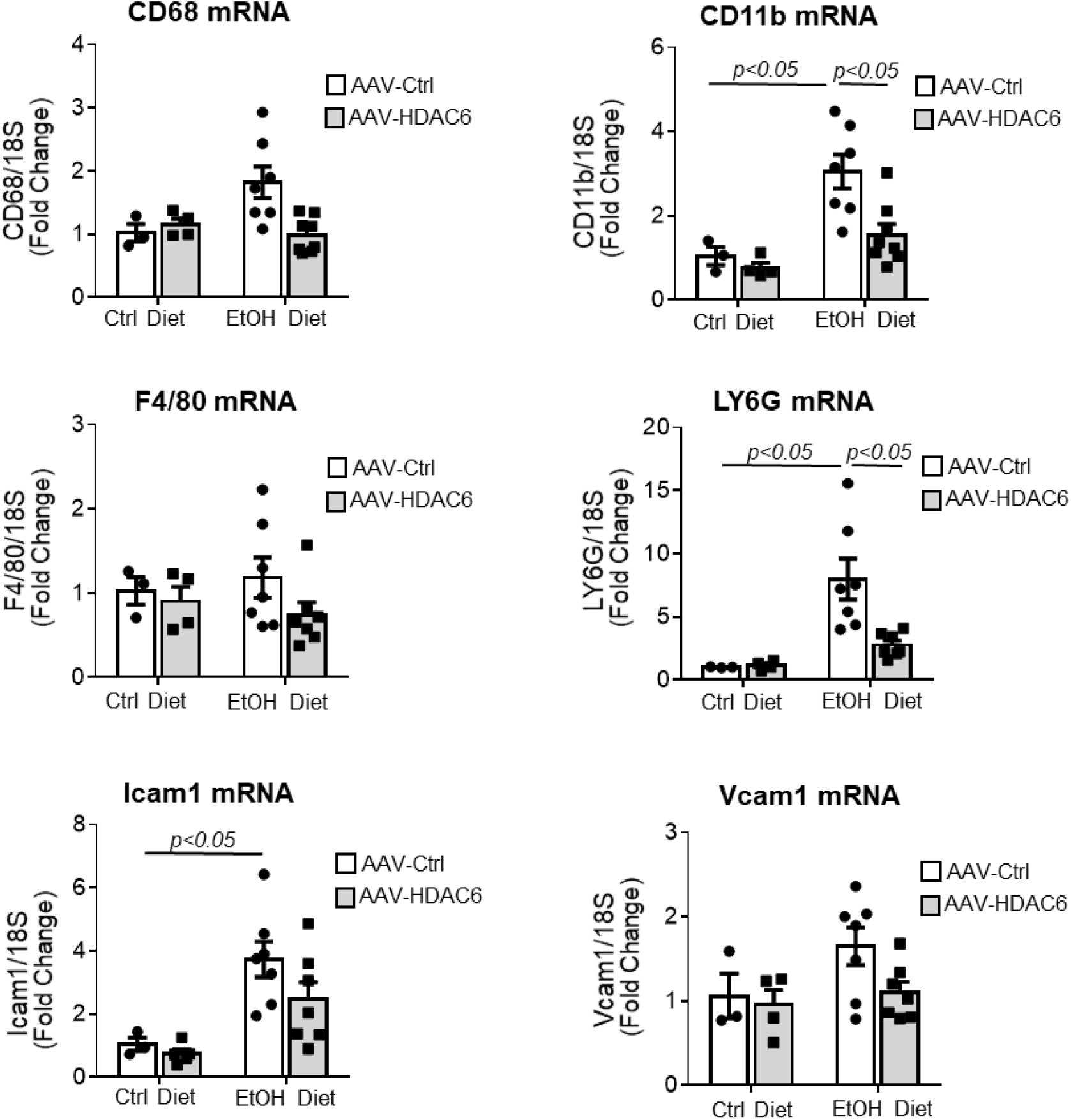
The effect of *in vivo* HDAC6 overexpression in liver ECs on alcohol-induced liver injury: mRNA expression. mRNA expression levels of genes related to inflammation, such as CD68, CD11b, F4/80, LY6G, Icam1 and Vcam1, in livers of Cdh5-CreERT2 mice injected with AAV-Ctrl or AAV-HDAC6 and fed with a control or an ethanol diet. n = 3 in AAV-Ctrl group and n = 4 in AAV-HDAC6 group for Control diet, n = 7 per group for Ethanol diet.

## Materials and Methods

### Acetyl-CoA assay

Acetyl-CoA contents in human LSECs were determined according to the manufacturer’s protocols (MAK039, Sigma-Aldrich, St. Louis, MO) using a fluorometric assay method. Briefly, cells were frozen rapidly and pulverized. Samples were deproteinized and precipitated by 1 N perchloric acid (34288, Sigma-Aldrich) while they were kept cold. The samples were then homogenized thoroughly and centrifuged at 13,000 xg for 10 minutes. Supernatants were neutralized with 3 M potassium bicarbonate solution (60339, Sigma-Aldrich). The samples were then cooled on ice for 5 minutes, and a pH was verified to be in the range of 6–8. The samples were spun for 2 minutes and supernatants were used. A reaction solution consisted of 20 μl sample solution, 21 μl acetyl-CoA assay buffer, 2 μl acetyl-CoA substrate mix, 1 μl conversion enzyme, 5 μl acetyl-CoA enzyme mix and 2 μl fluorescent probe, and was incubated at room temperature for 10 minutes. Fluorescence intensity was measured (λex = 535/ λem = 587 nm) and calculated using an acetyl-CoA standard curve.

### Serum ALT measurement

Serum ALT concentrations were measured using Liquid ALT Reagent Set (Pointe Scientific, Canton, MI). Reagent mix (R1:R2 = 5:1) was added into 96-well plates (100 μl per well) and pre-warmed at 37 °C for 5 minutes. Then, 10 μl of serum samples were transferred to each well and incubated at 37 °C for 1 minute. Absorbance at 340 nm was read by a plate reader every minute and recorded for 15 minutes. ALT levels were calculated by the mean absorbance difference/minute.

### Histological analysis

Liver tissues were fixed with formalin and embedded with paraffin. Paraffin sections were stained with hematoxylin and eosin (H&E) solution for histopathological analysis. All stained slides were scored for hepatic steatosis and lobular inflammation using a scoring system described by Kleiner DE et al (1). A total pathological score was determined by adding scores for steatosis, inflammation and liver cell injury. An experienced pathologist scored H&E-stained liver sections.

### CK8 immunohistochemistry and quantification

Immunohistochemical (IHC) staining was performed on paraffin-embedded liver specimens obtained from ethanol fed mice with AAV-Ctrl or AAV-HDAC6 using a VECTASTAIN ABC Kit (Vector Laboratories, Burlingame, CA) according to the manufacturer’s instructions. A primary antibody used was a monoclonal rat anti-mouse keratin 8 (K8) antibody (clone TROMA-I, 1:100, Developmental Studies Hybridoma Bank, The University of Iowa, Iowa City, IA). The number of CK8-negative hepatocytes in Zones 1 and 2 was counted. CK8 is a cytoskeletal intermediate filament protein in hepatocytes. It is known that CK8-negative hepatocytes are more susceptible to various forms of liver injury (2). IHC with CK8 has been used to assess ballooned hepatocytes in alcoholic hepatitis and nonalcoholic fatty liver disease (NASH) (3).

### Immunofluorescence

Liver tissues were fixed with paraformaldehyde (PFA) or formalin and then embedded. Frozen sections were washed with PBS for 10 minutes 3 times. Paraffin sections were de-paraffinized with xylene and dehydrated with graded ethanol. Antigen retrieval was performed using BD solution in a steamer for 15 minutes. The sections were then incubated with blocking buffer (5% donkey serum and 0.3% Triton X-100 in PBS) for 1 hour. After blocking, the sections were incubated with CYP2E1 (1:100, ab28146, Abcam, Cambridge, UK) and PECAM-1 (1:100, 550274 or 550300, BD Pharmingen, San Diego, CA) overnight at 4°C. After being washed with PBS 3 times, secondary antibodies (donkey anti-rabbit Alexa 647, 1:300; donkey anti-rat Alexa 647, 1:300; donkey anti-mouse Alexa 488, 1:300, Invitrogen, Waltham, MA) were applied for 30 minutes at room temperature. All samples were mounted with FluoroshieldTM containing DAPI (Sigma-Aldrich) and observed by a fluorescence microscope (Zeiss Observer Z1, Oberkochen, Germany) or a confocal microscope (Leica SP5, Wetzlar, Germany).

LSECs or hepatocytes were plated on coverslips and fixed in 4% PFA for 15 minutes. Then, cells were permeabilized with 0.1% Triton X-100 for 10 minutes and blocked by 5% donkey serum for 1 hour. After blocking, samples were incubated with CYP2E1 (1:100, ab28146, Abcam) overnight at 4°C. After washing, they were incubated with a secondary antibody (donkey anti-rabbit Alexa 488, 1:300, Invitrogen) for 30 minutes at room temperature. Then, the samples were mounted with FluoroshieldTM containing DAPI (Sigma-Aldrich) and observed under a fluorescence microscope (Zeiss Observer Z1). ER structure was visualized by staining with ER-Tracker™ Red dye (E34250, Molecular Probes, Eugene, OR) at a concentration of 1 μM for 15 minutes at 37°C. After washing, CYP2E1 proteins were stained following the above-mentioned procedure without fixation.

### Quantitative real-time polymerase chain reaction

Total RNA was extracted by TRIzol (Invitrogen) from LSECs or liver tissues according to the manufacturer’s instruction. RNA quantity and quality were assessed by micro-volume spectrophotometry on the NanoDrop 2000 (Thermo Fisher Scientific, Waltham, MA). Total RNA (1 μg) was reverse transcribed into cDNA using a Reverse Transcription Reagents kit (04897030001, Roche Molecular Systems, Branchburg, NJ). Quantitative real-time PCR (qPCR) was performed for cDNA samples using either TaqMan Universal Master Mix (Applied Biosystems, Foster City, CA) or PowerUp SYBR Green Master Mix (Applied Biosystems) on the ABI 7500 real-time PCR system (Applied Biosystems). 18S was used as a loading control. Results are shown as fold-changes relative to control groups. Primer specific sequences are listed in Supplementary Table 1.

### Western blot analysis

Cells or liver tissues were homogenized in a lysis buffer with protease inhibitor cocktail (Roche Diagnostics, Mannheim, Germany) and kept on ice for 30 minutes. Lysates were centrifuged at 12,000 rpm for 10 minutes at 4°C. Protein concentrations in the supernatants were determined using the BCA assay. An equal amount of protein (30-50 μg) from each sample was loaded onto sodium dodecyl sulfate–polyacrylamide gel electrophoresis gels and transferred to 0.22 μm polyvinylidene fluoride or nitrocellulose membranes. Membranes were probed with primary antibodies including CYP2E1 (1:800, ab28146, Abcam), Hsp90 (1:1000, 4877, Cell Signaling Technology, Danvers, MA), Hsp90 (1:3000, 610419, BD Biosciences, San Jose, CA), anti-Flag (1:1000, Agilent Technologies, Santa Clara, CA), eNOS (1:1,000, 610298, BD Biosciences), ACSS1 (1:1,000, sc-377149, Santa Cruz Biotechnology, Dallas, TX), ADH (1:1000, sc-133207, Santa Cruz Biotechnology), ALDH1/2 (1:1,000, sc-166362, Santa Cruz Biotechnology), HDAC6 (1:500, H2287, Sigma-Aldrich) and mouse anti–*β*-actin (1:1,000, A1978, Sigma-Aldrich). After washing, membranes were incubated with fluorophore-conjugated secondary antibodies. Detection and quantification of bands were performed using the Odyssey Infrared Imaging System (LI-COR Biosciences, Lincoln, NE). GAPDH or *β*-actin was used as a loading control.

### HUVEC ethanol experiment (For Supplementary Figure)

For ethanol treatment, human umbilical vein endothelial cells (HUVECs) were seeded on 6-well plates. After treatment with 100 mM ethanol in culture medium for 20 hours, an additional 100 mM ethanol was added to medium and incubated for 4 hours.

For acetaldehyde treatment, HUVECs were incubated with 50 μM acetaldehyde (402788, Sigma-Aldrich) in culture medium for 24 hours. An additional 50 μM acetaldehyde was added to medium at 4, 8 and 20 hours after incubation.

### Time-dependent NO assay (For Supplementary Figure)

HUVECs were seeded on 6-well plates. After 24 hours, medium was changed to phenol-red free culture medium (31053036, Gibco, San Diego, CA). Then, cells were treated with 100 mM ethanol at 0, 12, 16 and 20 hours after changing the medium. The media was collected 24 hours after the change to phenol-red free culture medium, resulting in 24, 12, 8 and 4-hour ethanol treatment, respectively, and processed for the measurement of nitrite, a stable end product of NO in aqueous solution. Nitrite concentrations were measured by the DAN assay.

---

**Supplementary Table 1.**
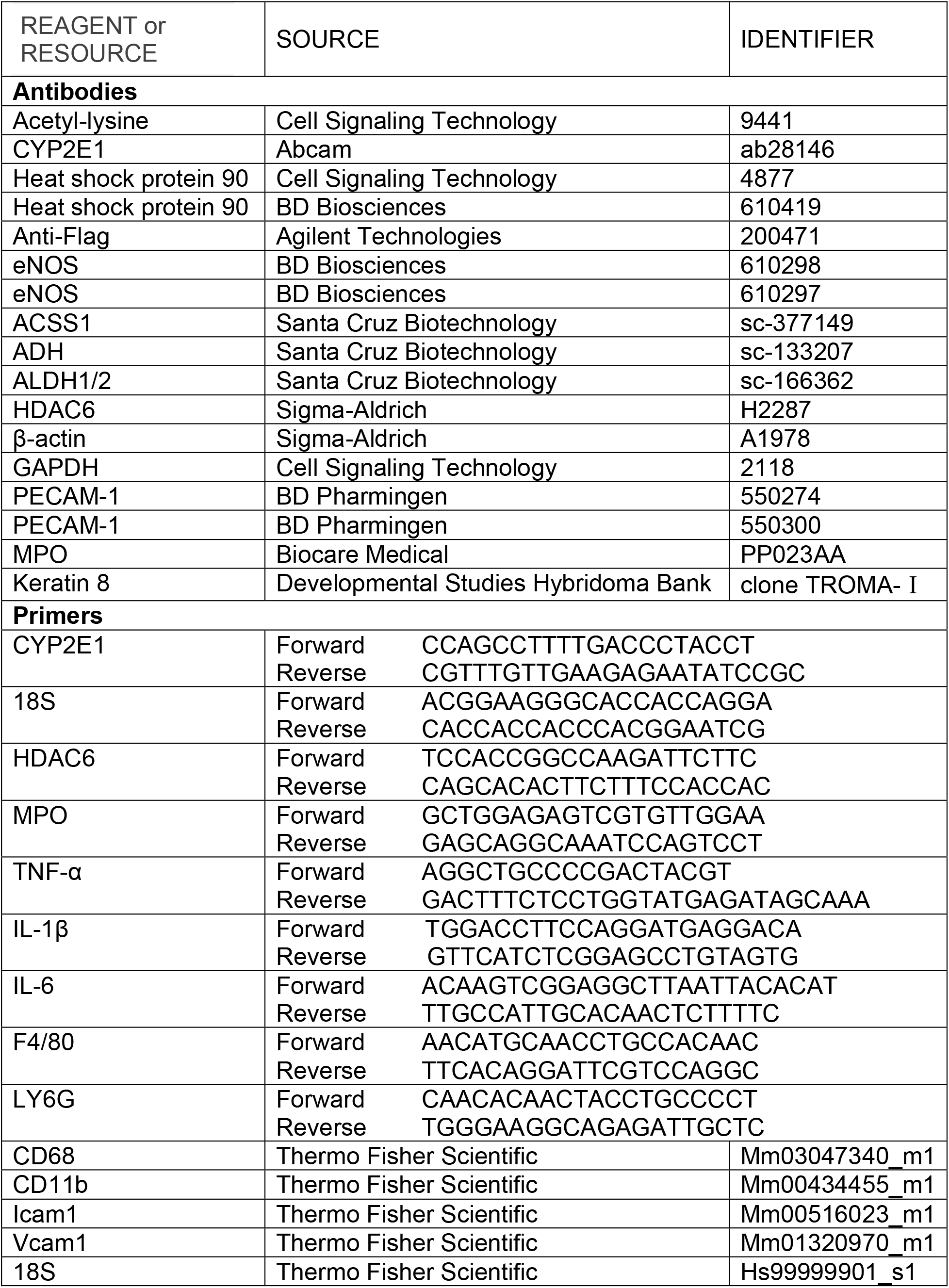
A list of antibodies and primers.

## References

1. Sorensen KK, Simon-Santamaria J, McCuskey RS, Smedsrod B. Liver Sinusoidal Endothelial Cells. Compr Physiol 2015;5:1751–1774.

2. Ruart M, Chavarria L, Camprecios G, Suarez-Herrera N, Montironi C, Guixe-Muntet S, Bosch J, et al. Impaired endothelial autophagy promotes liver fibrosis by aggravating the oxidative stress response during acute liver injury. J Hepatol 2019;70:458–469.

3. Pasarin M, La Mura V, Gracia-Sancho J, Garcia-Caldero H, Rodriguez-Vilarrupla A, Garcia-Pagan JC, Bosch J, et al. Sinusoidal endothelial dysfunction precedes inflammation and fibrosis in a model of NAFLD. PLoS One 2012;7:e32785.

4. McCuskey RS, Bethea NW, Wong J, McCuskey MK, Abril ER, Wang X, Ito Y, et al. Ethanol binging exacerbates sinusoidal endothelial and parenchymal injury elicited by acetaminophen. J Hepatol 2005;42:371–377.

5. Nanji AA, Tahan SR, Khwaja S, Yacoub LK, Sadrzadeh SM. Elevated plasma levels of hyaluronic acid indicate endothelial cell dysfunction in the initial stages of alcoholic liver disease in the rat. J Hepatol 1996;24:368–374.

6. Edenberg HJ. The genetics of alcohol metabolism: role of alcohol dehydrogenase and aldehyde dehydrogenase variants. Alcohol Res Health 2007;30:5–13.

7. Dey A, Cederbaum AI. Alcohol and oxidative liver injury. Hepatology 2006;43:S63–74.

8. Zakhari S. Overview: how is alcohol metabolized by the body? Alcohol Res Health 2006;29:245–254.

9. Novak RF, Woodcroft KJ. The alcohol-inducible form of cytochrome P450 (CYP 2E1): role in toxicology and regulation of expression. Arch Pharm Res 2000;23:267–282.

10. Koop DR, Chernosky A, Brass EP. Identification and induction of cytochrome P450 2E1 in rat Kupffer cells. J Pharmacol Exp Ther 1991;258:1072–1076.

11. Leung TM, Nieto N. CYP2E1 and oxidant stress in alcoholic and non-alcoholic fatty liver disease. J Hepatol 2013;58:395–398.

12. Schopf FH, Biebl MM, Buchner J. The HSP90 chaperone machinery. Nat Rev Mol Cell Biol 2017;18:345–360.

13. Scroggins BT, Robzyk K, Wang D, Marcu MG, Tsutsumi S, Beebe K, Cotter RJ, et al. An acetylation site in the middle domain of Hsp90 regulates chaperone function. Mol Cell 2007;25:151–159.

14. Garcia-Cardena G, Fan R, Shah V, Sorrentino R, Cirino G, Papapetropoulos A, Sessa WC. Dynamic activation of endothelial nitric oxide synthase by Hsp90. Nature 1998;392:821–824.

15. Gratton JP, Fontana J, O’Connor DS, Garcia-Cardena G, McCabe TJ, Sessa WC. Reconstitution of an endothelial nitric-oxide synthase (eNOS), hsp90, and caveolin-1 complex in vitro. Evidence that hsp90 facilitates calmodulin stimulated displacement of eNOS from caveolin-1. J Biol Chem 2000;275:22268–22272.

16. Galdieri L, Zhang T, Rogerson D, Lleshi R, Vancura A. Protein acetylation and acetyl coenzyme a metabolism in budding yeast. Eukaryot Cell 2014;13:1472–1483.

17. Boyer JL, Phillips JM, Graf J. Preparation and specific applications of isolated hepatocyte couplets. Methods Enzymol 1990;192:501–516.

18. Fulton D, Gratton JP, McCabe TJ, Fontana J, Fujio Y, Walsh K, Franke TF, et al. Regulation of endothelium-derived nitric oxide production by the protein kinase Akt. Nature 1999;399:597–601.

19. Nussler AK, Glanemann M, Schirmeier A, Liu L, Nussler NC. Fluorometric measurement of nitrite/nitrate by 2,3-diaminonaphthalene. Nat Protoc 2006;1:2223–2226.

20. Shah V, Haddad FG, Garcia-Cardena G, Frangos JA, Mennone A, Groszmann RJ, Sessa WC. Liver sinusoidal endothelial cells are responsible for nitric oxide modulation of resistance in the hepatic sinusoids. J Clin Invest 1997;100:2923–2930.

21. Iwakiri Y, Satoh A, Chatterjee S, Toomre DK, Chalouni CM, Fulton D, Groszmann RJ, et al. Nitric oxide synthase generates nitric oxide locally to regulate compartmentalized protein S-nitrosylation and protein trafficking. Proceedings of the National Academy of Sciences 2006;103:19777–19782.

22. Sangwung P, Greco TM, Wang Y, Ischiropoulos H, Sessa WC, Iwakiri Y. Proteomic identification of S-nitrosylated Golgi proteins: new insights into endothelial cell regulation by eNOS-derived NO. PLoS One 2012;7:e31564.

23. Valenzuela-Fernandez A, Cabrero JR, Serrador JM, Sanchez-Madrid F. HDAC6: a key regulator of cytoskeleton, cell migration and cell-cell interactions. Trends Cell Biol 2008;18:291–297.

24. Wang L, Wang H, Bell P, McCarter RJ, He J, Calcedo R, Vandenberghe LH, et al. Systematic evaluation of AAV vectors for liver directed gene transfer in murine models. Mol Ther 2010;18:118–125.

25. Wang BY, Ju XH, Fu BY, Zhang J, Cao YX. Effects of ethanol on liver sinusoidal endothelial cells-fenestrae of rats. Hepatobiliary Pancreat Dis Int 2005;4:422–426.

26. Nanji AA, Greenberg SS, Tahan SR, Fogt F, Loscalzo J, Sadrzadeh SM, Xie J, et al. Nitric oxide production in experimental alcoholic liver disease in the rat: role in protection from injury. Gastroenterology 1995;109:899–907.

27. Wang X, Abdel-Rahman AA. Effect of chronic ethanol administration on hepatic eNOS activity and its association with caveolin-1 and calmodulin in female rats. Am J Physiol Gastrointest Liver Physiol 2005;289:G579–585.

28. Shepard BD, Joseph RA, Kannarkat GT, Rutledge TM, Tuma DJ, Tuma PL. Alcohol-induced alterations in hepatic microtubule dynamics can be explained by impaired histone deacetylase 6 function. Hepatology 2008;48:1671–1679.

29. Lopez-Moreno JA, Marcos M, Calleja-Conde J, Echeverry-Alzate V, Buhler KM, Costa-Alba P, Bernardo E, et al. Histone Deacetylase Gene Expression Following Binge Alcohol Consumption in Rats and Humans. Alcohol Clin Exp Res 2015;39:1939–1950.

30. Ding G, Liu HD, Huang Q, Liang HX, Ding ZH, Liao ZJ, Huang G. HDAC6 promotes hepatocellular carcinoma progression by inhibiting P53 transcriptional activity. FEBS Lett 2013;587:880–886.

31. Qiu W, Wang B, Gao Y, Tian Y, Tian M, Chen Y, Xu L, et al. Targeting Histone Deacetylase 6 Reprograms Interleukin-17-Producing Helper T Cell Pathogenicity and Facilitates Immunotherapies for Hepatocellular Carcinoma. Hepatology 2019.

32. Jung KH, Noh JH, Kim JK, Eun JW, Bae HJ, Chang YG, Kim MG, et al. Histone deacetylase 6 functions as a tumor suppressor by activating c-Jun NH2-terminal kinase-mediated beclin 1-dependent autophagic cell death in liver cancer. Hepatology 2012;56:644–657.

33. Yin Z, Xu W, Xu H, Zheng J, Gu Y. Overexpression of HDAC6 suppresses tumor cell proliferation and metastasis by inhibition of the canonical Wnt/β-catenin signaling pathway in hepatocellular carcinoma. Oncol Lett 2018;16:7082–7090.

34. Yang HD, Kim HS, Kim SY, Na MJ, Yang G, Eun JW, Wang HJ, et al. HDAC6 Suppresses Let-7i-5p to Elicit TSP1/CD47-Mediated Anti-Tumorigenesis and Phagocytosis of Hepatocellular Carcinoma. Hepatology 2019;70:1262–1279.

35. Shepard BD, Tuma DJ, Tuma PL. Lysine acetylation induced by chronic ethanol consumption impairs dynamin-mediated clathrin-coated vesicle release. Hepatology 2012;55:1260–1270.

36. Kannarkat GT, Tuma DJ, Tuma PL. Microtubules are more stable and more highly acetylated in ethanol-treated hepatic cells. J Hepatol 2006;44:963–970.

37. Groebner JL, Giron-Bravo MT, Rothberg ML, Adhikari R, Tuma DJ, Tuma PL. Alcohol-induced microtubule acetylation leads to the accumulation of large, immobile lipid droplets. Am J Physiol Gastrointest Liver Physiol 2019;317:G373–G386.

38. Park PH, Miller R, Shukla SD. Acetylation of histone H3 at lysine 9 by ethanol in rat hepatocytes. Biochem Biophys Res Commun 2003;306:501–504.

39. Shukla SD, Restrepo R, Fish P, Lim RW, Ibdah JA. Different mechanisms for histone acetylation by ethanol and its metabolite acetate in rat primary hepatocytes. J Pharmacol Exp Ther 2015;354:18–23.

40. Taubert D, Berkels R, Grosser N, Schroder H, Grundemann D, Schomig E. Aspirin induces nitric oxide release from vascular endothelium: a novel mechanism of action. Br J Pharmacol 2004;143:159–165.

41. Arunachalam G, Yao H, Sundar IK, Caito S, Rahman I. SIRT1 regulates oxidant- and cigarette smoke-induced eNOS acetylation in endothelial cells: Role of resveratrol. Biochem Biophys Res Commun 2010;393:66–72.

42. Mattagajasingh I, Kim CS, Naqvi A, Yamamori T, Hoffman TA, Jung SB, DeRicco J, et al. SIRT1 promotes endothelium-dependent vascular relaxation by activating endothelial nitric oxide synthase. Proc Natl Acad Sci U S A 2007;104:14855–14860.

43. Jung SB, Kim CS, Naqvi A, Yamamori T, Mattagajasingh I, Hoffman TA, Cole MP, et al. Histone deacetylase 3 antagonizes aspirin-stimulated endothelial nitric oxide production by reversing aspirin-induced lysine acetylation of endothelial nitric oxide synthase. Circ Res 2010;107:877–887.

44. Farin FM, Pohlman TH, Omiecinski CJ. Expression of cytochrome P450s and microsomal epoxide hydrolase in primary cultures of human umbilical vein endothelial cells. Toxicol Appl Pharmacol 1994;124:1–9.

45. Haorah J, Knipe B, Leibhart J, Ghorpade A, Persidsky Y. Alcohol-induced oxidative stress in brain endothelial cells causes blood-brain barrier dysfunction. J Leukoc Biol 2005;78:1223–1232.

46. Ambade A, Mandrekar P. Oxidative stress and inflammation: essential partners in alcoholic liver disease. Int J Hepatol 2012;2012:853175.

47. Lu Y, Zhuge J, Wang X, Bai J, Cederbaum AI. Cytochrome P450 2E1 contributes to ethanol-induced fatty liver in mice. Hepatology 2008;47:1483–1494.

48. Butura A, Nilsson K, Morgan K, Morgan TR, French SW, Johansson I, Schuppe-Koistinen I, et al. The impact of CYP2E1 on the development of alcoholic liver disease as studied in a transgenic mouse model. J Hepatol 2009;50:572–583.

49. Cho YE, Mezey E, Hardwick JP, Salem NJr., Clemens DL, Song BJ. Increased ethanol-inducible cytochrome P450-2E1 and cytochrome P450 isoforms in exosomes of alcohol-exposed rodents and patients with alcoholism through oxidative and endoplasmic reticulum stress. Hepatol Commun 2017;1:675–690.

## References

1. Kleiner DE, Brunt EM, Van Natta M, Behling C, Contos MJ, Cummings OW, Ferrell LD, et al. Design and validation of a histological scoring system for nonalcoholic fatty liver disease. Hepatology 2005;41:1313–1321.

2. Ku NO, Strnad P, Zhong BH, Tao GZ, Omary MB. Keratins let liver live: Mutations predispose to liver disease and crosslinking generates Mallory-Denk bodies. Hepatology 2007;46:1639–1649.

3. Torruellas C, French SW, Medici V. Diagnosis of alcoholic liver disease. World J Gastroenterol 2014;20:11684–11699.

